# Two Contrasting Classes of Nucleolus-Associated Domains in Mouse Fibroblast Heterochromatin

**DOI:** 10.1101/484568

**Authors:** Anastassiia Vertii, Jianhong Ou, Jun Yu, Aimin Yan, Hervé Pagès, Haibo Liu, Lihua Julie Zhu, Paul D. Kaufman

**Affiliations:** Department of Molecular, Cellular and Cancer Biology, University of Massachusetts Medical School, Worcester, MA 01605 USA; Program in Computational Biology, Fred Hutchinson Cancer Research Center, Seattle, Washington, 98109-1024, USA

**Keywords:** nucleolus associated domains, NADs, chromatin territories, nucleolus, histone modification

## Abstract

In interphase eukaryotic cells, almost all heterochromatin is located adjacent to the nucleolus or to the nuclear lamina, thus defining Nucleolus-Associated Domains (NADs) and Lamina–Associated Domains (LADs), respectively. Here, we determined the first genome-scale map of murine NADs in mouse embryonic fibroblasts (MEFs) via deep sequencing of chromatin associated with purified nucleoli. We developed a Bioconductor package called *NADfinder* and demonstrated that it identifies NADs more accurately than other peak-calling tools, due to its critical feature of chromosome-level local baseline correction. We detected two distinct classes of NADs. Type I NADs associate frequently with both the nucleolar periphery and with the nuclear lamina, and generally display characteristics of constitutive heterochromatin, including late DNA replication, enrichment of H3K9me3 and little gene expression. In contrast, Type II NADs associate with nucleoli but do not overlap with LADs. Type II NADs tend to replicate earlier, display greater gene expression, and are more often enriched in H3K27me3 than Type I NADs. The nucleolar associations of both classes of NADs were confirmed via DNA-FISH, which also detected Type I but not Type II probes enriched at the nuclear lamina. Interestingly, Type II NADs are enriched in distinct gene classes, notably factors important for differentiation and development. In keeping with this, we observed that a Type II NAD is developmentally regulated, present in MEFs but not in undifferentiated embryonic stem (ES) cells.

## Introduction

In eukaryotic cell nuclei, genomic loci are distributed in three-dimensional space in a nonrandom manner. In particular, heterochromatic regions frequently reside at the nuclear lamina (NL) and/or in the perinucleolar region (van Koningsbruggen et al. 2010; Németh et al. 2010; Kind et al. 2013; Ragoczy et al. 2014; Kind et al. 2015). Loci clustered in these regions are generally less transcriptionally active and display histone modification patterns characteristic of poorly transcribed genes (reviewed in (Politz et al. 2013)). A major question in eukaryotic cell biology is how non-random localizations of loci are achieved and how they foster functional sequestration or, conversely, how interactions among accessible regions are co-regulated (Schoenfelder et al. 2009). Multiple studies suggest that positioning of human cell heterochromatin to either the lamina of perinucleolar region appears to be stochastic, and reshuffled at each mitosis when chromosomes are condensed and then decondensed in the subsequent interphase (van Koningsbruggen et al. 2010; Kind et al. 2013). Heterochromatin interactions are also regulated during cellular differentiation (Peric-Hupkes et al. 2010; Meuleman et al. 2013). These long-range chromosome interactions in mammalian cells are of great interest because they can regulate the developmental timing (Vernimmen et al. 2007) or the variegation of gene expression (Noordermeer et al. 2011). However, mechanisms governing these dynamics remain poorly understood.

As the site of ribosome biogenesis, nucleoli are organized around rDNA repeats (McStay and Grummt 2008; Pederson 2011) and consist of liquid-phase separated and functionally distinct layers of proteins and RNAs (Brangwynne et al. 2011; Feric et al. 2016). The perinucleolar chromatin that is the subject of this work surrounds nucleoli. This perinucleolar localization is a functionally relevant example of heterochromatin-mediated silencing, because transgene proximity to mammalian nucleoli is correlated with gene repression (Fedoriw et al. 2012b). An important step for analysis of the relevant silencing mechanisms is deep sequencing of nucleolar-associated DNA. To date, this has been analyzed in human cells and the plant *Arabidopsis thaliana* (van Koningsbruggen et al. 2010; Németh et al. 2010; Dillinger et al. 2017; Pontvianne et al. 2016), and these studies have detected specific nucleolar-associated domains (NADs). Consistent with their heterochromatic status, NADs are enriched in the H3K9me3 histone modification characteristic of constitutive heterochromatin, frequently include large multigene clusters, and are correlated with low gene expression and density.

Dramatic changes in the architecture of nucleolar-associated satellite DNA repeats during mouse development (Aguirre-Lavin et al. 2012) have been noted, but systematic characterization of murine NADs had not previously been done. To lay the groundwork for studying the biological roles of NADs in a system that allows analysis of mammalian developmental processes, we have mapped the nucleolar-associated domains (NADs) in mouse embryonic fibroblasts (MEFs). Also, previous studies of human NADs have differed regarding use of formaldehyde crosslinking and cell lines used (van Koningsbruggen et al. 2010; Németh et al. 2010; Dillinger et al. 2017). Notably, direct tests of the effects of crosslinking have not been performed. Therefore, we directly tested the effects of crosslinking, which we found had small effects on NAD identification, indicating that heterochromatin is naturally tightly associated with nucleoli. As expected, the majority of NADs are also LADs, and we refer to these here as Type I NADs. However, we detected a subset of mouse NADs that are not preferentially associated with the nuclear lamina. Therefore, these analyses have identified a distinct Type II subset of NADs. Compared to the Type I NADs, the Type II subset displays greater levels of gene expression and H3K27me3 modification, earlier DNA replication timing, and is enriched for particular classes of genes. Additionally, we show via DNA-FISH analyses that a Type II NAD in MEFs is indeed not enriched at the nucleolar periphery in ES cells, indicating regulation during differentiation. Therefore, the methods and data presented here demonstrate a subset of nucleolar heterochromatin that is not easily explained by stochastic positioning. Further, this work provides a platform for future investigation of the dynamics of the nucleolar heterochromatic compartment.

## Results

### Isolation of nucleoli from crosslinked and non-crosslinked MEFs

For our initial analyses of murine NADs, we compared nucleoli prepared with or without formaldehyde crosslinking (Fig. 1A; Supplemental Fig.1A). Both methods yielded highly-purified nucleoli based on phase microscopy images (Supplemental Fig. 1B; see also the “NAD-seq” QC files at the NIH 4DN Data Portal (data.4dnucleome.org)). Quantitative PCR suggested that rDNA sequences were enriched relative to the input total genomic DNA > 20-fold in the non-crosslinked preparations, and > 7-fold in crosslinked preparations (Supplemental Fig. 1C). Immunoblot analyses showed enrichment of nucleolar protein fibrillarin relative to non-nucleolar proteins such as actin, a nucleoporin (Nup62), or lamin A/C (Supplemental Fig.1D). Additionally, we characterized the small RNAs present in the non-crosslinked preparation, observing that the nucleoli were highly enriched for the nucleolar U3 RNA (Supplemental Fig.1E). Small RNAs could not be recovered from crosslinked samples (data not shown).

**Figure 1.**
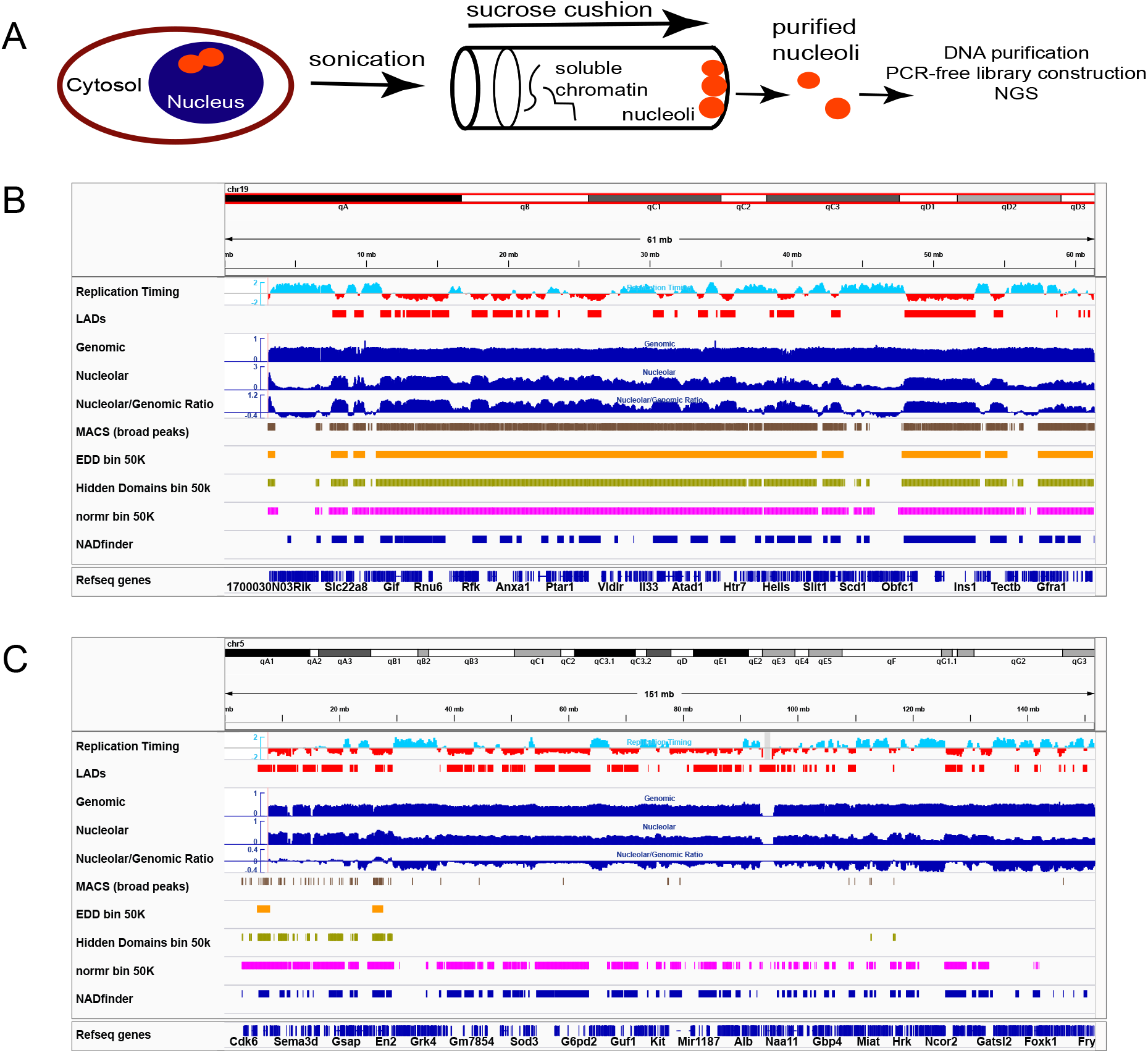
Identification of mouse nucleolus-associated domains (NADs). A. A schematic workflow of nucleolar isolation. Details that differ between the crosslinked and non-crosslinked versions of the preparation are illustrated in Supplemental Figure S1.
B. Comparison of NADs to heterochromatin features, and analysis of NAD-seq by multiple bioinformatic packages, including *NADfinder*. All data shown are from MEF cells. In this panel, the strongly nucleolar-associated chromosome 19 is shown in its entirety. Tracks shown from the top are DNA Replication timing data (Hiratani et al. 2010)(early replicating regions have positive values) and LADs (red, (Peric-Hupkes et al. 2010)). Below those are raw sequence data from crosslinked cells (blue, experiment #26): read counts for total genomic DNA (Genomic), nucleolar-associated DNA (Nucleolar), and the Nucleolar/Genomic ratio. Next are shown comparisons of peaks called by the bioinformatic packages MACS (broad peaks setting), EDD (50K bins), Hidden Domains (50K bins), and normr (50K bins, v1<0.01, q<10e-03). At the bottom are peaks called by *NADfinder* (default cutoff q<0.05). All software tested except *NADfinder* called almost the entire chromosome as peaks. NADs called by *NADfinder* generally correlate with LADs and late replication timing.
C. As in Figure 1B, showing chromosome 5. Note the gradual decrease in the ratio going from left to right (away from the centromere on the left of this acrocentric chromosome). Unlike MACs and EDD, NADfinder was able to call peaks on regions distal from the centromere.

### Bioinformatic analysis of nucleolar-associated DNA

We performed three biological replicate experiments for both the crosslinked and non-crosslinked protocols. For each experiment, we purified DNA from the isolated nucleoli, alongside genomic DNA from a sample of whole, unfractionated cells from the same population for normalization of the nucleolar sequencing data. These DNAs were used to generate sequencing libraries via PCR-free protocols, yielding genome-scale maps. Visual inspection of these “NAD-seq” data showed that the total genomic DNA was largely evenly distributed across the genome (Fig. 1B, C), occasionally interspersed with short, poorly represented regions. The nucleolar DNA was distributed with distinct peaks and valleys. The ratio of nucleolar/genomic read counts provides a raw metric of enrichment of nucleolar association.

We expected mouse nucleolar-associated regions to be highly enriched in heterochromatin, as observed in human cells (van Koningsbruggen et al. 2010; Németh et al. 2010; Dillinger et al. 2017). Therefore, we compared our raw data to datasets that distinguish mouse euchromatin and heterochromatin. For example, late DNA replication (Hiratani et al. 2008; 2010) and association with the nuclear lamina (Peric-Hupkes et al. 2010) are properties generally associated with heterochromatin (these features are colored red in Figs. 1B-C). Notably, peaks in our raw nucleolar-associated DNA sequence data frequently overlapped these features. Additionally, we observed that the patterns of nucleolar enrichment often appeared similar in each of the triplicate experiments performed with the crosslinked (XL) and non-crosslinked (nonXL) protocols (Supplemental Fig. S2). Consistent with those profiles, Pearson correlation analysis of the nucleolar/genomic ratios from each of the six experiments displayed a high degree of similarity between crosslinked and non-crosslinked data (Supplemental Figure S2). Together, these data suggested that both of our biochemical preparation methods reproducibly yield mouse nucleolar heterochromatin.

To facilitate comparisons to other genomic features, we then sought to call peaks in our datasets, and tested several established bioinformatic packages developed for analysis of LADs or ChIP-seq data (Fig. 1B-C, Supplemental Fig. 3). We tested MACS, which is frequently used for the analysis of ChIP-Seq data (Zhang et al. 2008; Feng et al. 2012). Because of the large size of heterochromatin regions like NADs and LADs compared to most transcription factor binding sites, we also tested EDD (Lund et al. 2014), hidden domains (Starmer and Magnuson 2016), and normr (Kinkley et al. 2016; Helmuth et al. 2016), packages suited for detecting enrichments over large genomic regions. We tested each of these using at least two different settings (Fig. 1 and Supplemental Fig. 3). Small chromosomes, which are more frequently associated with nucleoli (Quinodoz et al. 2018), were often annotated as almost entirely nucleolus associated using these tools (e.g. chr19, Fig. 1B), without distinguishing most peaks and valleys observed in the nucleolar DNA sequencing data. Conversely, we observed that for longer chromosomes, most of these packages identified very few peaks on the chromosome end distal from the centromere (pictured on the right side of these acrocentric mouse chromosomes, e.g. chr5 in Fig. 1C). We noted that the nucleolar/genomic ratios displayed a notable slope across large chromosomes, with higher average ratio values near the centromere-containing end of the acrocentric mouse chromosomes. This likely reflects the frequent nucleolar association of pericentric heterochromatin (Ragoczy et al. 2014). However, even in regions far from the centromeres, there are still local peaks in the nucleolar/genomic ratio, often coinciding with heterochromatic features. These observations suggested that normalizing data to a local background would facilitate accurate identification of peaks in regions that are not near centromeres on larger chromosomes, while also facilitating discrimination between peaks and valleys on small chromosomes.

Therefore, we developed a Bioconductor package termed *NADfinder* to analyze our data that includes local baseline correction (Supplemental Figure 4A-C and **Material and Methods**). We note that local baseline correction was required for the *NADfinder* program to identify peaks distal to the centromere. That is, without local baseline correction, only few strong peaks near the centromere-proximal end of the chromosome were identified (green tracks, Supplemental Fig. 4B, C), reminiscent of the stronger centromere-proximal peaks in the raw data (Fig. 1C). In contrast, inclusion of the baseline correction step generated peak profiles that are distributed across the chromosomes (blue tracks, Supplemental Fig. 4B, C). Notably, these baseline-corrected peaks frequently overlap with lamin-associated regions and late replicating regions (red, Supplemental Fig. 4B, C), consistent with our expectation that NADs are largely heterochromatic.

In the first biological replicate experiment for each purification method, we sequenced approximately 200 million reads. To determine whether this amount was necessary, we performed preliminary peak calling using *NADfinder*, and then performed subsampling analyses (Supplemental Fig. 4D, E). Subsampling analysis results showed that the % of peaks detected and overlapped with those from the whole dataset initially increases quickly with increased depth of sequencing, and plateaus at a subsampling depth of approximately 25% (i.e. 50 million reads). We therefore performed 50 million reads for samples in the subsequent second and third replicate experiments for both experimental protocols (crosslinked (“XL”) and noncrosslinked (“nonXL”)). We designed *NADfinder* with functionality for analyzing datasets with biological replicates and for generating adjusted p-values for each peak. We used this feature to generate the preliminary “NADfinder” tracks shown in Fig. 1 and Supplemental Fig. 3.

### Distinct subsets of nucleolar-associated domains

In addition to MEF LADs (Guelen et al. 2008; Peric-Hupkes et al. 2010), we analyzed the “constitutive interLADs” (ciLADs), which are sequences that are not lamin-associated at any stage during neural differentiation of mouse embryonic stem cells, first into neural precursor cells and then astrorocytes (Peric-Hupkes et al. 2010; Meuleman et al. 2013). That is, ciLADs generally represent euchromatic regions that display much higher levels of gene activity than do LAD regions. To analyze the overlap among NADs, LADs and ciLADs, we separated each nucleotide of our NAD peaks into subsets of those overlapping with one of the other datasets, or neither. For example (chr17, Fig. 2A), NADs (blue bars) and LADs (red) displayed frequent regions of overlap (magenta), in both the XL and nonXL datasets. Analysis of the overlaps in a genome-wide fashion (Fig. 3B), indicated that the majority but not all NAD peak sequences overlap with LADs, regardless of the nucleolus purification method. For example, ciLADs (Fig. 2A, cyan) overlap with a subset of NADs (green). The ciLAD-overlapped subset was smaller than the LAD-associated regions, but occurred in similar regions in samples from the two experimental protocols (XL and nonXL). Thus, we had subdivided NADs into LAD-overlapped and ciLAD-overlapped subsets. For convenience, we will refer to these as Type I and Type II, respectively. Additionally, a subset of NADs overlapped with neither LADs nor ciLADs (Fig. 2B).

**Figure 2.**
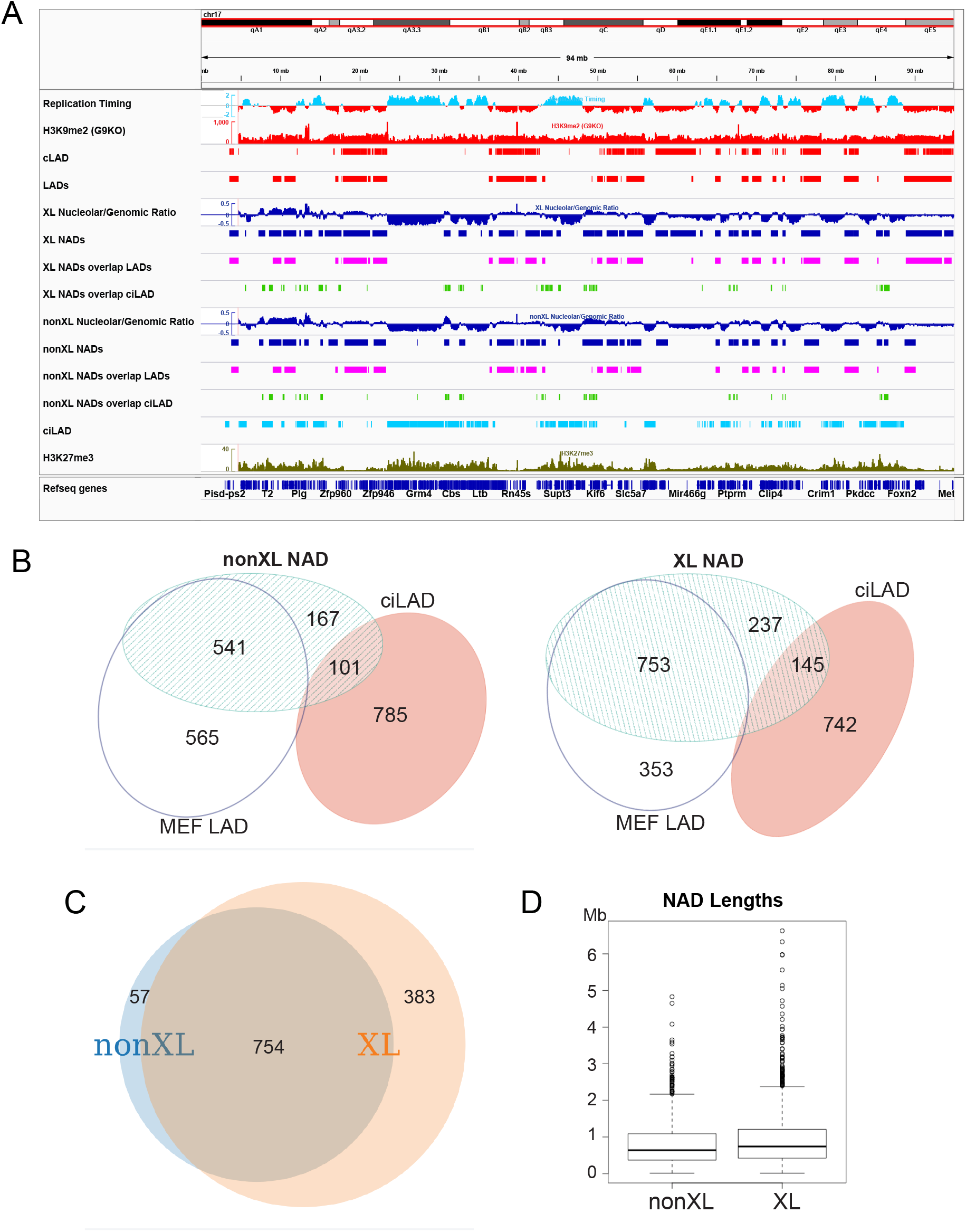
Distinct classes of mouse NADs. A. Subdivision of MEF NADs based on overlap with LAD or ciLAD regions. chr17 is shown. Features associated with heterochromatin (H3K9me2, lamin association, and late replication timing) are colored in red, while features associated with euchromatin (early replication timing and ciLAD regions) are in cyan. At the top is shown DNA replication timing (Hiratani et al. 2010), followed by H3K9me2 (from MEF cells lacking methyltransferase G9a, making heterochromatic H3K9me2 more easily visualized (GSM2803100)(Chen et al. 2018)). Next are constitutive LAD and MEF LAD peaks (Peric-Hupkes et al. 2010; Meuleman et al. 2013). Following those are eight tracks of NAD data, the top four from a crosslinked preparation (XL experiment #26), and the bottom four from a non-crosslinked preparation (nonXL experiment #29). At the top of each set is the log2 ratio of nucleolar/genomic sequencing reads, followed by the peaks determined by *NADfinder* (blue). These are followed by a pair of tracks in which the NAD peaks were separated at the nucleotide level into regions that overlap the MEF LADs (magenta) and regions that overlap ciLADs (light green). The bottom two tracks are ciLADs in cyan (Peric-Hupkes et al. 2010; Meuleman et al. 2013), and H3K27me3 in olive green ((GSM2417097)(Chronis et al. 2017)).
B. Venn diagram illustrating the overlap among NADs, LADs and ciLAD regions. Data from non-crosslinked experiments are on the left, and crosslinked ones are on the right. Numbers show the size of the indicated regions, in Mb.
C. Venn diagram illustrating overlap of XL and nonXL NAD peaks. Numbers indicate size of each region in Mb, which are drawn proportional to size.
D. Boxplot distributions of NAD lengths, showing median and quartiles in the box.

**Figure 3.**
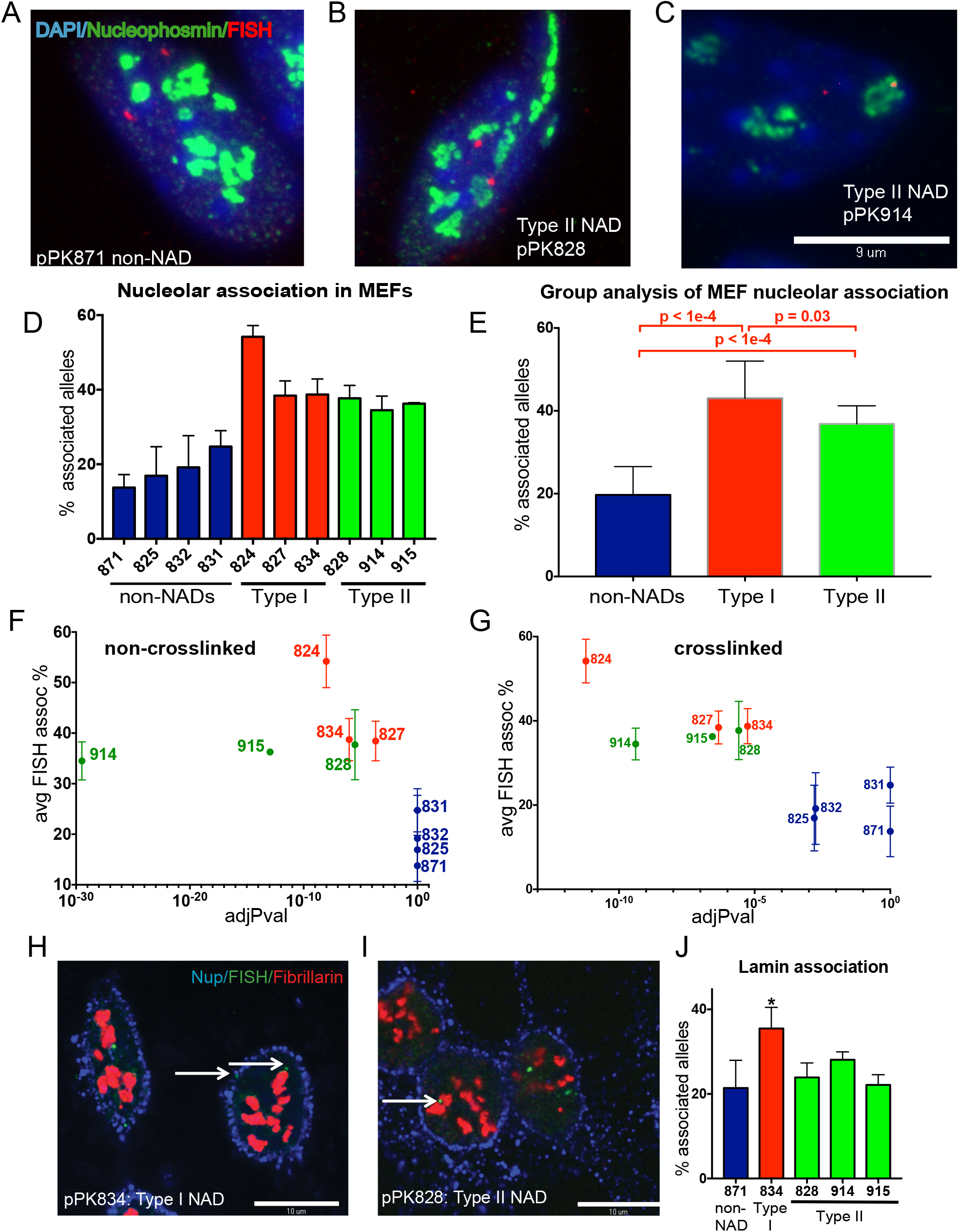
3-D DNA-FISH experiments confirm nucleolar association of NADs. A. Maximum projection of confocal microscopy images from a 3-D immunoFISH experiment (pPK871 BAC DNA probe, red; fibrillarin, green; DAPI, blue; scale bar 10μm).
B. As in panel A, with pPK828 as the DNA probe.
C. As in panel A, with pPK914 as the DNA probe.
D. Graph of percentage of alleles that are associated with nucleolus (mean ± SEM for n ≥ 3 biological replicate experiments). Negative probes (blue) display nucleolar association that is not significantly different than the negative probe pPK871 (p > 0.05, Welch’s t-test). In contrast, all type I (red) and type II (green) probes display association significantly different from pPK871 (p < 0.03).
E. Data for nucleolar association from panel D were grouped (see coloration). p values for the differences between the different groups are shown.
F. Graph comparing the % nucleolar-associated alleles from DNA-FISH (y-axis) vs the adjusted p-value (x-axis) of the nonXL NAD peaks identified by *NADfinder* that overlap with the indicated BAC probe. Note that the “negative” probes (blue) have an adj.p-value = 1 = 10e0.
G. As in panel F except that x-axis is the adjusted p-value of the XL NAD peaks. Note that the “negative” probes (blue) have an adj.p-value > 10e-3.
H. Single optical sections of confocal microscopy images of a 3-D immunoFISH experiment. pPK834 BAC DNA probe hybridizing to a type I NAD, green; fibrillarin, red; nuclear pore protein Nup62, blue; scale bar 10 μm.
I. As in panel C, with pPK828 as the DNA probe hybridizing a type II NAD.
J. Graph of percentage of alleles that are associated with nuclear periphery (mean ± SEM for n ≥ 3 biological replicate experiments). Only Type I NAD probe pPK834 (*) displays laminar association that is significantly different than the negative control pPK871 (p = J. 042, Welch’s t-test). As expected, three Type II NAD probes (pPK828, 914 and 915) do not display significant laminar association.

We next compared NADs in our XL and nonXL datasets. Genome-scale analysis showed that almost all peaks in the nonXL dataset were also found in the XL dataset. Additionally, crosslinking resulted in a greater proportion of the genome identified as NAD peaks (Fig. 2C). Specifically, the NAD peaks comprised a total of 1137 Mb in the XL and 811 Mb in the nonXL datasets, representing 41% and 30% of the non-repetitive mouse genome, respectively. For comparison, LADs comprise approximately 40% of the mouse genome (Guelen et al. 2008; Peric-Hupkes et al. 2010), and NADs in human diploid IMR90 fibroblasts cover 38% of genome (Dillinger et al. 2017). Also, XL and nonXL murine NADs had similar median lengths (Fig. 2D), comparable to MEF (534kb) and human fibroblast LADs (553kb) (Guelen et al. 2008; Peric-Hupkes et al. 2010). Therefore, the overall characteristics of mouse NADs are similar to those of other large heterochromatic domains. However, as shown below, these genome averages mask distinct subsets within the population of mouse NADs.

### Immuno-DNA FISH validation of genomic data

To validate peaks called by *NADfinder*, we performed DNA fluorescent in situ hybridization (DNA-FISH) experiments. We used bacterial artificial chromosome (BAC) probes to detect specific genomic loci, and anti-fibrillarin or anti-nucleophosmin antibodies to visualize nucleoli. We performed at least three biological replicate experiments per probe, scoring the percentage of alleles (n > 100) in a population of cells associated with nucleoli (Fig. 3A-C). First, we chose a negative control BAC (termed pPK871) that does not contain nucleolus-enriched sequences in either the crosslinked or non-crosslinked data (Supplemental Fig. 5A). As expected, we observed a low frequency of overlap between pPK871 and nucleoli (median value of 14% of alleles scored, Fig. 3D), similar to the values observed for non-nucleolar associated probes in human cell studies (van Koningsbruggen et al. 2010; Németh et al. 2010; Smith et al. 2014; Dillinger et al. 2017). We analyzed multiple additional probes to characterize NAD associations in MEFs (e.g. Figs. 3B, C; Supplemental Figure S5), and observed that probes from both “Type I” and “Type II” NAD peaks displayed significantly greater frequencies of association with nucleoli than did pPK871. This was evident in the measurement of the percentage of total alleles associated for the individual probes analyzed (Fig. 3D), and when data for the probes of each group were combined (Fig. 3E). Additionally, we compared the observed percentage of nucleolar association of FISH probes to the adjusted p-value (q value) of the corresponding peaks calculated by *NADfinder* (Fig. 3F-G). For both the nonXL and XL datasets, we observed high q values for “negative” probes which displayed nucleolar association levels statistically indistinguishable from those of negative control pPK871 (Fig. 3F-G, colored in blue). In contrast, the Type I (green) and II (red) NAD probes displayed lower q values, clearly distinct from the negative probes (note that the x-axis in these graphs is logarithmic). We conclude that *NADfinder* identified bona fide NADs.

### Testing association of NADs with the nuclear lamina

Designation of Type II NADs was based on overlap with ciLAD regions, which do not associate with the nuclear lamina in MEFs or during neural differentiation of ES cells (Peric-Hupkes et al. 2010; Meuleman et al. 2013). However, we considered the possibility that some regions could have been classified as type II NADs if their apparent lack of laminar association was a false negative. We therefore directly tested a subset of our BAC probes for associations with the nuclear lamina in additional immunoDNA-FISH experiments. In this case, we tested for colocalization with the edge of DAPI-stained material, which can be highlighted by staining with antibodies against a nuclear pore protein (Fig. 3H-I). We observed that our negative control BAC, pPK871, displayed on average ~20% association with the lamina, whereas a Type I probe that overlaps with a LAD, pPK834, displays significantly higher association frequencies (Fig. 3J). In contrast, three different Type II probes displayed frequencies of lamina association that were not significantly different from those of the negative control. These data indicate that Type II NADs are a subset of heterochromatin that preferentially associates with nucleoli rather than the nuclear lamina in MEF cells.

### Biological properties of the NAD subsets

To explore NAD biological properties, we first compared gene densities of different NAD subsets to the rest of the genome (Fig. 4A). Consistent with their heterochromatic character, the complete sets of NADs identified by either the XL and nonXL protocols (red bars) display much lower average gene density than the genome average (brown). Furthermore, LAD-overlapping Type I NADs (pink bars), display even lower gene density levels. In contrast, the ciLAD-overlapped Type II NADs (purple bars) display greater gene densities than do Type I NADs, NADs that overlap with neither LADs nor ciLADs, or NADs as a whole. Next, we examined steady-state RNA levels in the different classes, expressed as FPKM in RNAseq data (Fig. 4B). We observed that NADs are much less frequent sites of gene expression than the genome-wide average. However, the Type II NADs are distinct in that they display elevated gene expression compared to the other subclasses, consistent with the euchromatic character of the ciLAD subset of the genome (Meuleman et al. 2013). In summary, these data indicate that NADs are not solely heterochromatic.

**Figure 4.**
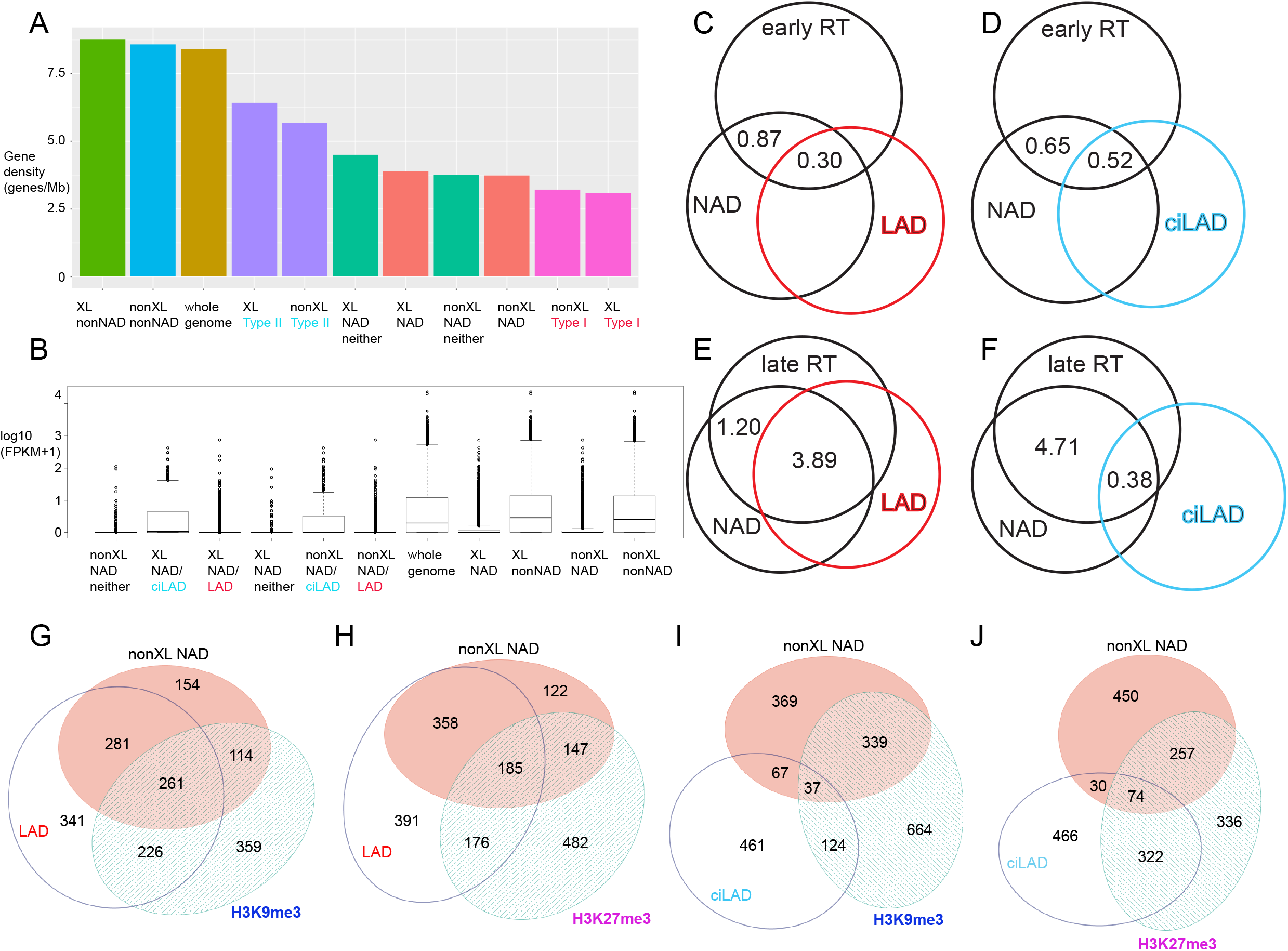
Differential chromatin states and gene density and expression in different classes of NADs. A. Gene densities of the indicated genomic regions are shown, ranked left to right.
B. Distribution of gene expression levels from RNAseq data, expressed as FPKM for the same genomic regions analyzed in panel A.
C. Venn diagram analysis of the overlap among NADs (nonXL), LADs and H3K9me3 peaks. Numbers indicate size of each region in Mb, which are drawn proportional to size.
D. As in panel C, except that the overlaps with LADs and H3K27me3 peaks are shown.
E. As in panel C, except that the overlaps with ciLADs and H3K9me3 are shown.
F. As in panel C, except that the overlaps with ciLADs and H3K27me3 are shown.
G. Venn diagram analysis of overlaps with DNA replication timing (RT) data and LADs. As in panels C-F, nonXL NAD data are analyzed, except that in place of histone modifications, overlaps with regions of early Rt are indicated. For simplicity, only the numbers (Mb) for the overlap between the NADs and the RT data are shown. The numbers here are smaller than those in panels C-F because the RT data is based on short microarray probes rather than deep sequencing.
H. As in panel G, except that overlaps with ciLADs and early RT are shown.
I. As in panel G, except that overlaps with LADs and late RT are shown.
J. As in panel G, except that overlaps with ciLADs and late RT are shown.

We next examined DNA replication timing (Fig. 4C-F), a genomic feature that largely distinguishes euchromatin and heterochromatin (Hiratani et al. 2008; 2010). We focused on the regions of overlap between the replication timing (RT) data and the NAD peaks (numbered regions in Fig. 4C-F). We note that these are representative of the genome-wide trends, but the absolute values are smaller than other overlaps we analyzed (e.g. Fig. 2B), because the RT data peaks consist of short microarray probes. We observed that NAD regions overlapping with early RT were more likely to also overlap with ciLADs (0.52 Mb) than LADs (0.30 Mb; Fig. 4C-D). A much more skewed distribution was observed for the late RT-NAD overlapped regions, which displayed a ten-fold larger overlap with LADs than ciLADs (Fig. 4E-F). Therefore, the LAD-overlapped Type I NADs are almost always late replicating; Type II NADs are distinct with their tendency towards early replication.

We then analyzed histone modifications related to heterochromatin, namely H3K9me3, a hallmark of constitutive heterochromatin that is generally inaccessible to transcription factors, and H3K27me3, a hallmark of facultative heterochromatin that is frequently regulated in a tissue-specific manner and is accessible to transcription factors and paused RNA polymerase (Becker et al. 2016). We observed that just under half of NAD regions overlapping with LADs (Type I NADs) also overlapped with H3K9me3 peaks (261 Mb of 542 Mb, Fig. 4G), while a smaller proportion (185 of 542 Mb) overlapped with H3K27me3 peaks (Fig. 4H). These data are consistent with the extensive H3K9me3 marking across LADs (Sadaie et al. 2013), which also feature less broad H3K27me3 peaks at their boundaries (Guelen et al. 2008; Harr et al. 2015). A more dramatic difference was observed upon inspection of the NAD-ciLAD overlapped regions (Type II NADs) (Fig. 4I-J). Only 37 Mb out of these 134 Mb overlapped H3K9me3 peaks (Fig. 4I), but twice as much (74 Mb) overlapped H3K27me3 peaks (Fig. 4J). Thus, compared to Type I NADs, Type II NADs lack lamin association, display greater gene density, expression, and H3K27me3 marks, and are more often early replicating. Thus, the Type II subset of NADs shares many features with facultative heterochromatin.

### Roles of H3K27 methylation and phase separation

A hallmark of facultative heterochromatin is methylation of histone H3K37 by the Ezh2 subunit of the Polycomb Repressive Complex 2 (PRC2, (Margueron and Reinberg 2011)). Previous studies implicated H3K27me3 in the localization of LADs to the nuclear periphery (Harr et al. 2015), so we tested whether the same would be true for nucleolar localization. Using two commercially available inhibitors of the Ezh2 methyltransferase, we observed that we could efficiently reduce cellular H3K27me3 levels, detected either via immunofluorescence or immunoblotting (Fig. 5A-E). Both these compounds significantly reduced both the nucleolar and the laminar association of Type I BAC probe, pPK834 (Figure 5F-J). Analysis of a Type II probe (pPK828) showed significant loss of nucleolar association in the presence of EPZ6438; the inhibitor GSK126 reduced association but not to a statistically significant level (Fig. 5K-O). pPK828 does not associate above background levels with the nuclear lamina, and neither drug altered that pattern. Together, these data indicate that H3K27 methylation is important for both nucleolar and laminar heterochromatic localizations.

**Figure 5.**
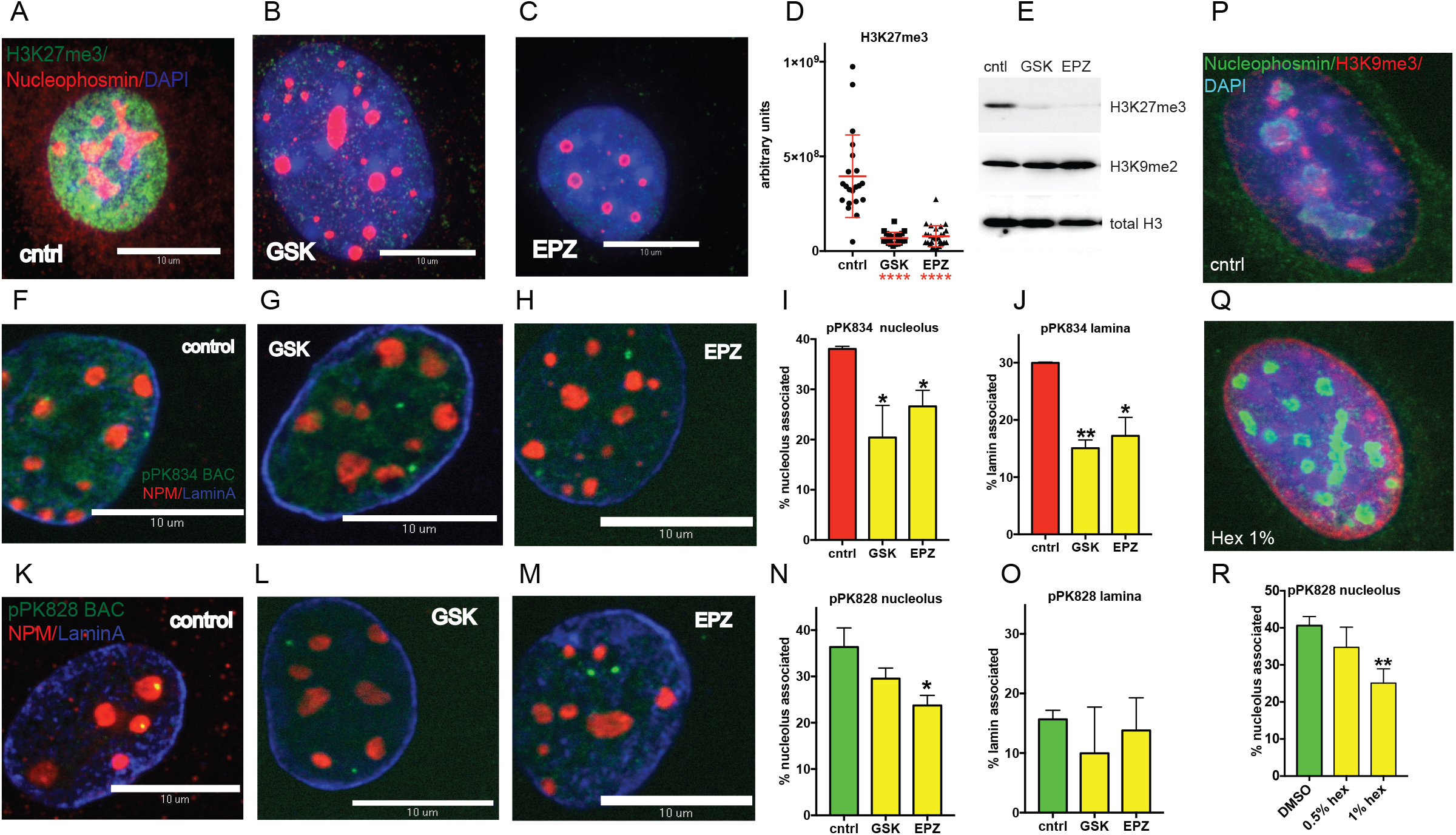
Inhibition of H3K27 methylation blocks NAD localization. A. Immunofluorescence measurement of H3K27me3 levels in control cells.
B. As in panel K, except cells were treated with GSK126.
C. As in panel K, except cells were treated with EPZ6438.
D. Quantitation of IF data from panels A-C. GSK126, ****: p < 0.0001, Welch’s t-test. EPZ6438, ****: p < 0.0001, Welch’s t-test.
E. Immunoblot analyses of cells treated as in panels A-C.
F. 3D immuno-FISH localization of type I probe pPK834 (green), anti-nucleophosmin antibodies (NPM (ab10530, Abcam), red), and anti-LaminA antibodies (blue). Untreated control cells.
G. As in Panel F, except that cells were treated with GSK126.
H. As above, except that cells were treated with EPZ6438.
I. Quantitation of pPK834 nucleolar association measured by DNA FISH as in Figure 3. GSK126, *: p = 0.040, Welch’s t-test. EPZ6438, *: p = 0.023, Welch’s t-test.
J. Quantitation of pPK834 laminar association measured by DNA FISH as in Figure 3. GSK126, **: p = 0.0029, Welch’s t-test. EPZ6438, *: p = 0.021, Welch’s t-test.
K. 3D immuno-FISH localization of type II probe pPK828 (green), with other markers as in Panel A. Untreated control cells.
L. As in Panel K, except that cells were treated with GSK126.
M. As above, except that cells were treated with EPZ6438.
N. Quantitation of pPK828 nucleolar association. GSK126, p = 0.083, Welch’s t-test. EPZ6438, *: p = 0.018, Welch’s t-test.
O. Quantitation of pPK828 laminar association. GSK126, p = 0.33, Welch’s t-test. EPZ6438, *: p = 0.63, Welch’s t-test.
P. Immunofluorescence analysis of H3K9me3 distribution in untreated control cells.
Q. As in panel P, treated with media containing 1% 1,6-hexanediol for 2 hours.
R. Quantification of pPK828 nucleolar association in control and hexanediol-treated cells. 1% hex **: p = 0.0090, Welch’s t-test.

Recent data indicate that higher order chromosomal interactions can be driven by phase separation, involving liquid demixing of heterochromatin components (Strom et al. 2017; Larson et al. 2017). Phase separation also contributes to the self-assembly of nucleoli themselves (Feric et al. 2016). One diagnostic used to study these compartmentalized interactions is the sensitivity of these to the aliphatic alcohol 1,6-hexandiol, which is thought to perturb liquid-like condensates via disruption of weak hydrophobic interactions (Ribbeck and Görlich 2002). We therefore tested whether features of MEF heterochromatin would also be sensitive to this compound. First, we observed that the pattern of H3K9me3 localization, which prominently decorates punctate foci of heterochromatin in untreated MEF cells (Guenatri et al. 2004; Michal Sobecki et al. 2016), was dramatically relocalized upon incubation of cells in media containing 1% hexanediol for two hours (Fig. 5P-Q). These data are consistent with large-scale heterochromatin alterations. We also noted that nucleoli themselves were still present, and that the 1% hexanediol concentration tested is below that used to disrupt other liquid demixed entities, such as the nematode synaptonemal complex (Rog et al. 2017), Drosophila HP1 clusters (Strom et al. 2017), yeast or human cell stress granules (Kroschwald et al. 2015; 2017), or nuclear pore complexes (Ribbeck and Görlich 2002). We therefore tested whether NAD interactions with nucleoli were affected by hexanediol after two hours of treatment. We observed a dose-dependent response, with a significant loss of nucleolar association of NADs at 1% hexanediol (Fig. 5R). These data suggest that NADs are a phase-separated form of heterochromatin.

### Type II NADs house developmentally regulated genes

In order to learn more about the potential for gene expression from type II NADs, we tested for enrichment of GO-terms among the genes in these regions (Fig. 6A-C; Supplemental Tables 2-3). Within the “Biological Processes” classifications, we observed the lowest adjusted p-values are for terms associated with developmental processes, for example organ morphogenesis and sensory organ development. Individual genes within these categories included the developmental transcription factors Pax2 (Fig. 6D) and Sox1 (Fig. S6A), and developmental signaling molecules such as Fgf1 (Fig. S6B) and Bmp4. Most notable among the “Cell Components” GO terms were keratin intermediate filament proteins, which can occur in gene clusters within a NAD (Fig. S6C). In addition, multiple types of ion channels were detected such as Scn sodium channel genes (Fig. S6C). The most significant GO-derived “Molecular Functions” classes included transcription factors; other classes include solute transporters such as glutamate transporter Grik3 (Fig. 6E). In summary, these data indicate that a distinct subset of transcription factors related to differentiation, and hallmarks of differentiated cell lineages including ion channels, are enriched within the Type II NADs in MEF cells.

**Figure 6.**
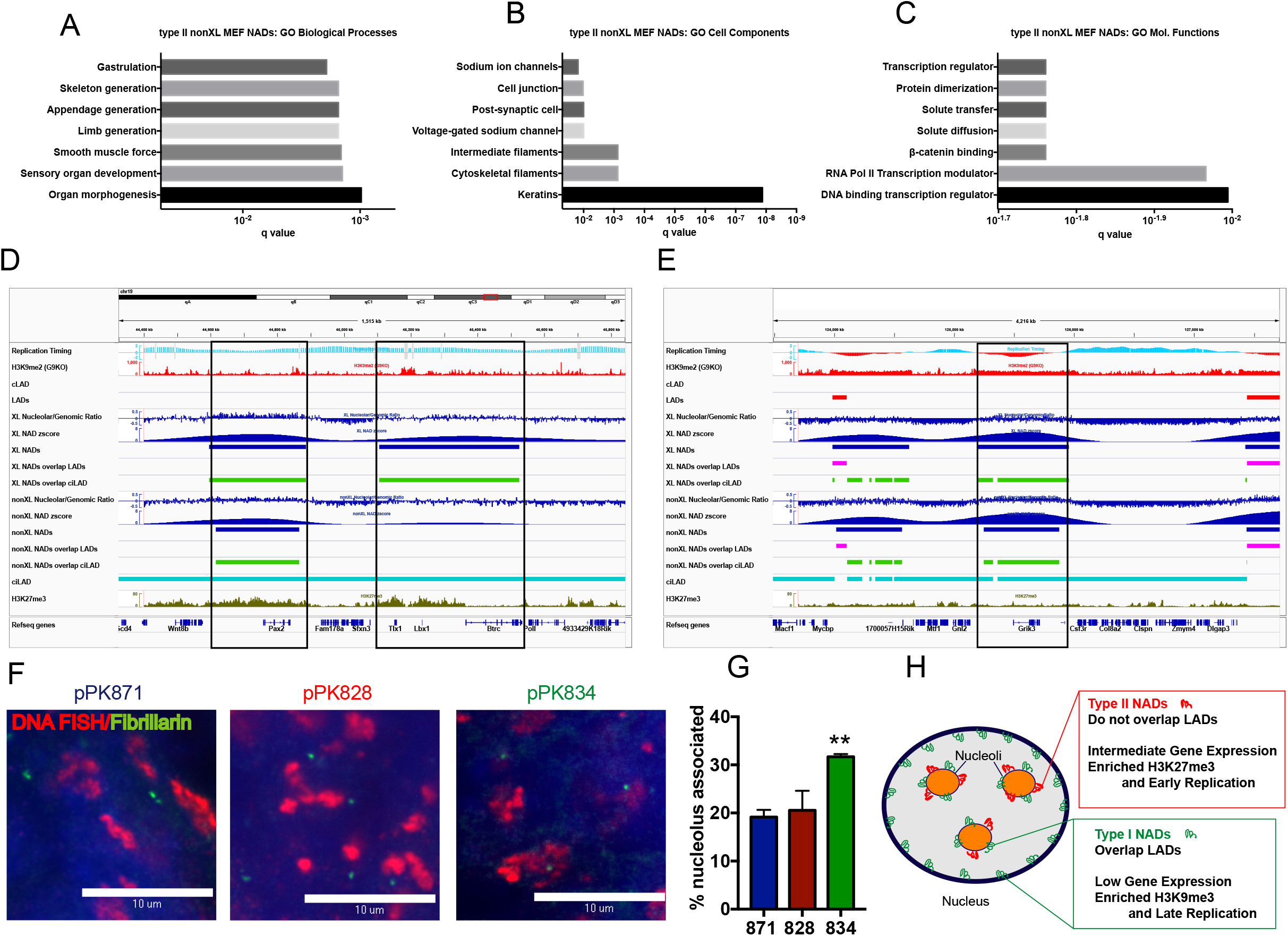
Type II NADs contain developmentally regulated genes and can be differentially localized in MEF and ES cells. A. GO enrichment analysis of type II NADs (nonXL). In this panel, the 7 most significant Biological Processes annotations and q values are shown.
B. As in panel A, showing the Cell Components annotations.
C. As in panel A, showing the Molecular Functions annotations.
D. Analysis of specific loci, using the same IGV session as in Figure 2, except that tracks showing *NADfinder-derived* zscores are also included. Genomic region showing two Type II NAD peaks (the one on the right achieved a statistical significance of q<10e-3 only in the XL but not nonXL data). The peak on the left contains the Pax2 transcription factor gene. The peak on the right contains the Tlx1 and Lbx1 homeobox transcription factor genes.
E. As in panel D, showing a Type II peak containing the Grik3 glutamate receptor gene.
F. Single optical sections of confocal microscopy images of 3-D FISH analysis of mouse D3 embryonic stem cells. BAC DNA probes are labeled in red, fibrillarin in green, scale bar 10μm. In MEFs, pPK871 is a negative control probe, pPK828 is a Type II NAD, and pPK834 is a Type I NAD (see Fig. 3).
G. Quantification of D3 mES cell FISH. Graph, left: percentage of alleles from the indicated probes associated with nucleoli (mean ± SEM). Welch’s t-tests for difference from pPK871: ** pPK834 (p = 0.0018), pPK828 (p = 0.62).
H. A diagram that summarizes the distinct characteristics of Type I and II NADs.

The observation that Type II NADs contain many developmentally regulated genes raised the question of whether there is developmental regulation of perinucleolar heterochromatic associations. As a first test of this idea, we performed DNA-FISH experiments using D3 mouse embryonic stem cells (Hiratani et al. 2010)(Fig. 6F-H). As we observed in MEF cells, the negative control BAC pPK871 displayed low levels of nucleolar association, and Type I probe pPK834 was significantly more frequently associated. In contrast, Type II probe pPK828, which was associated with nucleoli but not lamina in MEF cells (Figures 2-3), was not significantly nucleolar associated in D3 cells. These data indicate that NADs can adopt different localizations in different cell types.

## Discussion

### NADfinder, a new Bioconductor package for large domain analysis

In previous studies of NAD sequences (Németh et al. 2010; van Koningsbruggen et al. 2010; Dillinger et al. 2017; Pontvianne et al. 2016), bioinformatics tools for analyzing these datasets had not been developed. We found that tools developed for analyzing LADs and ChIP-seq data were suboptimal for the analysis of the mouse NADs. Specifically, multiple existing software packages failed to detect NADs in the regions distal to the centromere in long chromosomes and/or would label almost the entire short chromosomes as NADs in our MEF dataset (Figs. 1 and S3). We therefore developed a Bioconductor package called *NADfinder*. We observed that the use of local baseline correction enhanced recognition of peaks in the distal regions of large chromosomes, while improving the distinction of peaks on small chromosomes (Fig. S4). Importantly, peaks called by *NADfinder* were validated via DNA-FISH experiments (Fig. 3). We imagine that *NADfinder* will not only be useful for NAD studies, but will be globally useful for the analysis of large chromosome domains, particularly in cases where local background signals will differ significantly across the genome.

One advantage of deploying our software in the Bioconductor framework is that we can leverage the rich genomic annotation data and borrow functionalities already implemented in other R/Bioconductor packages. For example, we used the *ATACseqQC* package (Ou et al. 2018a) to efficiently read in the alignments, the *baseline* package for baseline correction, the *signal* package for smoothing, the *trackViewer* package (Ou et al. 2018b) for visualizing the signals, the *limma* package (Ritchie et al. 2015) for statistical analysis of the signal noise ratios across multiple replicates, and the *ChIPpeakAnno* package (Zhu et al. 2010; Zhu 2013) for annotating NADs with other genomic features of interest and for GO enrichment analyses. *NADfinder* is freely available at https://bioconductor.org/packages/release/bioc/html/NADfinder.html. Major functions are described in the Material and Method section. More detailed information on the commands and instructions for running the software are included in the associated user guide at https://bioconductor.org/packages/release/bioc/vignettes/NADfinder/inst/doc/NADfinder.html and https://bioconductor.org/packages/release/bioc/manuals/NADfinder/man/NADfinder.pdf.

### Diverse classes of perinucleolar heterochromatin

Previous studies in human cells suggested that there should be extensive overlap among LAD and NAD regions of the genome. Specifically, several experiments revealed extensive reshuffling of the interphase localizations of heterochromatin during each mitosis. For example, in an interphase cell, photoactivation of caged fluorescent histones near the nucleolar periphery results in the labeled molecules being distributed both at the nucleolar periphery and at the lamina in the two daughter cells after mitosis (van Koningsbruggen et al. 2010). Likewise, imaging of a GFP-DamID reader domain protein showed that LADs redistribute not only to lamina but also to nucleoli in daughter cells (Kind et al. 2013). These experiments indicated that the interphase localization of heterochromatin is largely stochastic, with apparent random placement of heterochromatic loci either at the lamina or surrounding nucleoli. Therefore, the expectation was that LADs and NADs would be largely the same.

Previous mapping of nucleolar heterochromatin has been done in human cells and in the plant Arabidopsis (Németh et al. 2010; van Koningsbruggen et al. 2010; Dillinger et al. 2017; Pontvianne et al. 2016). In human cells, the pioneering studies first describing perinucleolar heterochromatin (Németh et al. 2010; van Koningsbruggen et al. 2010) had low sequencing depth by today’s standards, but generally supported the idea that NADs are heterochromatic, with reduced gene density and expression relative to the genomic average. More recently, analysis of human IMR90 fibroblast heterochromatin emphasized the overlap and similarity between NADs and LADs, sharing enrichment for H3K9me3, gene and repeat density (Dillinger et al. 2017). In summary, the cell biological and genomic experiments mentioned above indicated that the majority of mammalian heterochromatin is directed in a stochastic manner to either the lamina or the perinculeolar region.

However, there have also been experiments suggesting that there are trans-acting factors that can specifically regulate particular heterochromatin destinations. For example, the lamin B receptor, an integral protein of the inner nuclear membrane, contributes to the laminar association of heterochromatin (Solovei et al. 2013). Conversely, the Chromatin Assembly Factor-1 p150 subunit governs the nucleolar localization of proliferation antigen Ki-67, which in turn governs NAD localization to nucleoli (Smith et al. 2014; Matheson and Kaufman 2017). The specificities observed in these studies suggest that not all heterochromatic loci will localize stochastically, and that some recruitment to specific locations can occur.

We also note that a recent study showed that an Ezh2-containing ribonucleoprotein complex termed MICEE tethers specific loci to the perinucleolar region (Singh et al. 2018). However, Ezh2 appears to have multiple roles, because inhibition of H3K27me3 inhibits both laminar {Harr:2015gj and nucleolar heterochromatin localizations (Fig. 5). Indeed, the latter data suggest that H3K27me3 could have a crucial and general role in heterochromatin localization, perhaps via contributing to phase separation (Fig. 5). However, there are likely to be multiple mechanisms, because the PRC2 complex that catalyzes H3K27me3 is not required for nucleolar localization of the Kcnq1 locus (Fedoriw et al. 2012a).

In summary, our data indicate that a significant subset of nucleolar heterochromatin is not frequently distributed to the lamina. Specifically, through locus-unbiased biochemical purification of nucleoli based on their sedimentation properties and resistance to sonication, we found distinct classes of loci in the associated DNA. The majority of NADs identified (Fig. 2B) were the expected loci that overlap with LADs, which we term Type I NADs here. However, we also identify Type II NADs, which were defined as peaks in our data that overlap with regions previously classified as “ciLAD”, that is, regions that do not associate with lamina at multiple stages of cellular differentiation (Peric-Hupkes et al. 2010; Meuleman et al. 2013). We hypothesize that Type II NADs will be enriched in sites of regulated localization and expression during differentiation, and we are now testing these ideas.

We note other recent studies have shown that sedimentation behavior of short crosslinked chromatin fragments helps detect functionally distinct subpopulations that cannot be distinguished by histone modification patterns alone (Becker et al. 2017). Therefore, we believe that biochemical studies can serve as an important alternative source of data for analyzing the structure and function of mammalian genomes.

## Materials and Methods

### Cell culture and treatments

Mouse embryonic fibroblasts (MEFs) derived from C57/Bl6 mice were immortalized using a standard 3T3 procedure (Todaro and Green 1963) at the UMASS Transgenic Mouse Modeling Core. Cells were obtained at p48. Cells were cultured in Dulbecco’s modified Eagle medium, DMEM (Life technologies, cat#11995-065), 10% bovine fetal serum, and 100 U/ml penicillin/ streptavidin. For passaging, old medium was aspirated, and cells were washed with 5 ml PBS per 10 cm plate. 1 ml of 0.05% (w/v) trypsin-EDTA was added and the plate was incubated at 37°C until the cells detached. 5 ml 37°C medium was added per 10 cm plate to quench the trypsin and the cells were transferred to a 15-ml conical tube and pelleted by centrifugation for 5 min at 200g at 4°C. Media was aspirated and cells were resuspended in 5 ml of fresh pre-warmed (37°C) medium. Cells were passaged at dilutions ranging from 1:2 to 1:10.

MEF cells were cloned by serial dilution. Cells diluted into single cell per well in 96-well plates, and wells containing healthy single clones were further expanded. Of these, Clone 5 cells were used in all experiments except uncrosslinked nucleolus isolation experiment #18 (see Supplemental Table S1), which used the uncloned population.

For Ezh2 inhibitor experiments, MEF cells were either treated with 0.1% DMSO or with the following inhibitors for 72 hours.

| Inhibitor | Cat. No. | Stock | Final conc |
| --- | --- | --- | --- |
| GSK126 | ApexBio #A3446 | 10 mM in DMSO | 10 μM |
| EPZ6438 | ApexBio #A8221 | 10 mM in DMSO | 100 nM |

1,6-Hexanediol (Sigma-Aldrich, #240117-50G) was dissolved directly into culture media (10% stock). MEFs were treated with a final concentration of 1 % or 0.5% 1,6-Hexanediol for 2 hrs.

Mouse embryonic stem cell (mES) line D3 was a kind gift from David Gilbert’s lab. The cells were grown in 2i mES medium on plates without gelatin coating, so that the cells grew in suspension. Below are components used for 2i mES medium:

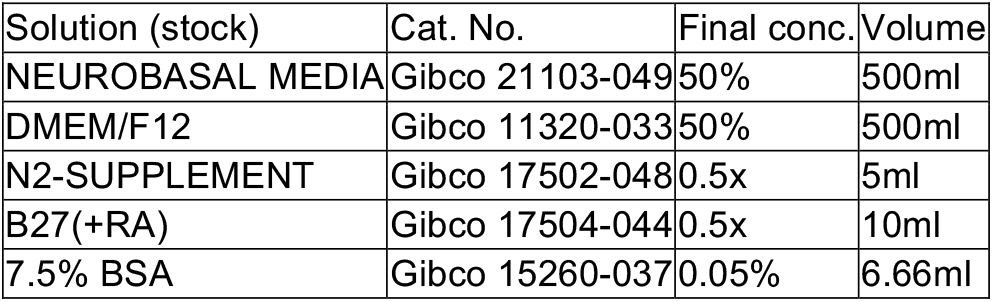
Serum free ES medium (SFES) - 1000 ml.

Complete “2i” medium - 100 ml.

Complete "2i" medium - 100 ml
| Solution (stock) | Cat. No. | Final conc. | Volume |
| --- | --- | --- | --- |
| SFES | N/A | 1x | 100ml |
| PD03259010 (10mM) | Reprocell 04-0006-02C | 1 μM | 10 μl |
| CHIR99021 (10mM) | Reprocell 04-0004-02C | 3 μM | 30 μl |
| GLUTAMINE (200mM) | Gibco 25030-081 | 2mM | 1ml |
| Monothioglycerol (11.9M) | Sigma M6145-25ML | 1.5x10<sup>-4</sup>M | 1.26 μl |
| LIF (1x10<sup>6</sup> U/ml)* | Cell Guidance Systems GFM200 | 1,000U/ml | 100 μl* |

Medium was kept out of the light at 4 °C, for no more than 2 weeks.

Cells were passaged by removing “2i” media and applying 0.5 ml of accutase (EMD Millipore SF006) per 3 cm plate for a few minutes at RT to separate cell colonies and to obtain a singlecell suspension. After incubation with accutase, 5 mL of SFES was added. The suspension was transferred into 15 ml tube and centrifuged at 1,000 rpm for 5 min at RT. The cell pellet was resuspended in complete “2i” media and plated into new vessel at a dilution of 1:6-1:10.

### MEF Nucleoli isolation

For each preparation, cells were grown in ten 15 cm plates until they reached 80-90% confluence, for a total of 30-100 x 10^6^ cells. Old media was removed and fresh media was added 1 h prior to nucleoli preparation. An additional plate grown in parallel was reserved for total genomic DNA extraction (Quick-DNA Universal Kit (Zymo Research, CA)).

### Non-crosslinked method of nucleoli isolation

The method was adapted from the Lamond lab protocol (van Koningsbruggen et al. 2010). Each dish was rinsed twice with 5 ml pre-warmed PBS, 3 ml 0.25% trypsin-EDTA were added per dish, followed by incubation at 37°C until cells were detached. 10 ml of complete media was added to each plate to quench the trypsin, and the cells were centrifuged in 50 ml conical tubes at 925 x rpm (200 x g) for 5 min (Sorvall Legend RT centrifuge) at 4°C. Cell pellets were washed with PBS and the packed cell volumes (PCV) were estimated. Five pellet volumes of PBS were added, cells were centrifuged and resuspended in ice-cold PBS, cell numbers were measured using a haemocytometer. At this point, the cells for the total genomic DNA sample were frozen. For the remainder of the cells, ~ 3 x PCV of ice-cold Nuclear Extract Buffer A (10 mM Hepes-KOH, pH 7.9, 10 mM KCl, 1.5 mM MgCl_2_,), plus 0.5 mM DTT was added freshly. Cells were incubated on ice for 10 min to promote swelling in this hypotonic buffer. Swelling was monitored using a Evos microscope with a 20X objective, using the phase contrast filter. Swollen cells were transferred by pouring into a glass Wheaton Dounce homogenizer (15-ml size, pre-rinsed with Nuclear Extract Buffer A) with a Type B Pestle, pre-chilled on ice. Cells were homogenized with 30 slow strokes, and lysis was assessed by examination of a small aliquot on the Evos microscope. The homogenization was stopped when >90% of the cells were burst, leaving intact nuclei and with various amounts of cytoplasmic material attached. 0.1-0.5 ml was saved as a total extract sample. Homogenized cells were centrifuged at 2000 rpm for 5 min at 4°C. A 0.5 ml aliquot of the supernatant was saved as “cytosolic extract”. The crude nuclear pellet was resuspended with 3 ml solution S1 (0.25 M sucrose, 10 mM MgCl_2_) and freshly added plus 0.5 mM DTT. Any pellet that could not be resuspended readily by gentle pipetting likely contained lysed nuclei and was discarded. Each resuspended pellet was layered over 4 ml of S2 (0.35 M sucrose, 0.5 mM MgCl_2_) solution in a 15-ml conical tube and centrifuged at 2000 rpm for 5 min at 4°C to enrich the purity of the nuclei. The purified nuclear pellets were then resuspended with 3 ml of S2 solution by gentle pipetting. A 0.1 ml sample of these resuspended nuclei was reserved as the “Nuclear” fraction. The nuclear suspension was sonicated with 10 second bursts (with 10 second intervals between each burst) using a Soniprep 150 (MSE) with a fine probe sonicator and set at power setting 5. The sonication was monitored using a phase contrast microscope. The sample was sonicated until almost no intact cells were detected and released nucleoli were observed as dense, refractile bodies. 0.7 ml of the sonicated sample was layered over 0.7 ml of S3 solution (0.88 M sucrose, 0.5 mM MgCl_2_) in 1.5 ml microfuge tubes and centrifuged at 20 seconds at 16,000 x g at 4°C. The pellet contains the nucleoli, and the supernatant (combining both the sucrose layer and the upper aqueous material) was retained as the “nucleoplasmic fraction”. The nucleoli were resuspended with 1.0 ml of S2 solution, combined in one microfuge tube, followed by centrifugation at 1430 x g for 5 min at 4°C. This wash was repeated twice. The pellet contains highly purified nucleoli. The nucleoli were resuspended in 0.2 ml of S2 solution, snap frozen in liquid nitrogen and stored at –80 °C.

### Crosslinked method of nucleoli isolation

The preparation of formaldehyde-crosslinked cell extracts was adapted from previous studies (Németh et al. 2010; Sullivan et al. 2001). Cells were grown as above and fixed with 1% formaldehyde added directly to the media and incubated at RT. After 10 min, the media was removed and the crosslinking was quenched by adding 5 ml 1M glycine. The cells were washed with ice-cold PBS, collected by scraping in 40 ml PBS, and centrifuged at 200 x g for 5 min at 4°C. The cell pellet was re-suspended in 1 ml high magnesium (HM) buffer (10 mM HEPES-KCl pH7.5, 0.88 M sucrose, 12 mM MgCl_2_, 1 mM DTT). Cells were then sonicated on ice (2-3 bursts for 10s at full power) using a Soniprep 150 (MSE) with a fine probe. The release of nucleoli was monitored microscopically. Nucleoli were pelleted by centrifugation in a microfuge at 15000g for 20s and re-suspended in 0.5 ml low magnesium (LM) buffer (10 mM HEPES-KCl pH 7.5, 0.88 M sucrose, 1 mM MgCl_2_, 1 mM DTT). The sample was subjected to a second round of sonication (1 burst 10s full power), and centrifuged at 15,000g for 20s to pellet nucleoli. The pellet was resuspended in 20/2 TE buffer (20 mM Tris-Cl pH 7.5, 2 mM EDTA) for immediate use or in 20/2TE + 50% glycerol to snap freeze in liquid nitrogen and store at −80°C.

### Quantitative PCR

DNA extraction from input whole cells and purified nucleoli was performed using Quick-DNA Universal Kit (Zymo Research, CA). DNA concentration was measured by Qubit dsDNA HS Assay kit (Invitrogen, Eugene, OR). 5 ng of DNA from each sample was analyzed using the Kappa Sybr Fast Q-PCR Kit (Kappa Biosystems, Wilmington, MA). The following primers were used: Set1 Fw: gag gtt gaa ggt ggt ttc ca; Rv: gag cag tcg ggt gct ctt ac; Set 2 Fw: gaa ctt tga agg ccg aag tg; Rv: atc tga acc cga ctc cct tt. The following program was used: 98°C 30s, 95°C 10s, 60°C 30s, 40 cycles. All the signals were normalized to that of genomic DNA as indicated in the figure legends, and the 2^-ΔΔ^CT method was used for quantification (Life Technologies).

### Antibodies

The following antibodies were used: fibrillarin (ab5821, Abcam, Cambridge, MA), B23 (nucleophosmin) (sc-32256, Santa Cruz, Dallas, TX), Nup62 (ab50008, Abcam Cambridge, MA), Lamin A/C (2032, Cell Signaling technology, Danvers, MA USA), Lamin A (ab25300, Abcam), Nucleophosmin (ab10530, Abcam), Histone H3 (di methyl K9) (ab1220, Abcam), Histone H3 K9me3 (ab8889, Abcam), H3 K27me3 (39155, ActifMotif), Histone H3 (ab18521, Abcam), actin (sc-8432, Santa Cruz, Dallas, TX). Secondary antibodies for immunofluorescence were conjugated with: Alexa 488, Cy3 (Jackson ImmunoResearch, West Grove, PA). For western blots, horseradish peroxidase (HRP) anti-mouse and anti-rabbit secondary antibodies (Jackson ImmunoResearch, West Grove, PA) were used.

### Immunoblotting

Protein concentrations from total cell lysates or isolated nucleoli were assessed by Bradford assay (BioRad reagent Blue R-250, cat. #161-0436 with BSA as a standard). 10 μg of each sample were loaded per lane on 12% SDS-PAGE gels, transferred to PVDF membrane for 2hrs, 80v, 4°C. Membrane was blocked in PBS-nonfat milk and incubated with corresponding antibodies in accordance to manufacture instructions.

### Small RNA isolation

Small RNAs were isolated using Qiagen miRNAeasy kits. In brief, we prepared RNA from 500 μl from each fraction that was collected during non-crosslinked nucleoli preparations. 700 μl QIazol reagent was added, and the mixture was vortexed and incubated at RT for 5 min. After incubation, 140 μl of chloroform was added, vortexed, incubated for 2 min at RT and clarified by centrifugation for 15 min at 12,000g at 4° C. The aqueous clear phase was transferred in to new tube and 1.5x volume of 100% ethanol was added, mixed by pipetting, and loaded onto the RNAeasy column. Non-bound material was removed by centrifugation, and the column was washed with RPE buffer and eluted in 50 μl of RNAse-free water. 3 μl of RNAse inhibitor (Promega, N2511) was added to each sample. 1 μg of RNA from total, cytosolic, nuclear, nucleoplasmic and nucleolar fractions analyzed on a 10% denaturing 7 M urea acrylamide gel and stained with SybrGreen (Sigma-Aldrich, #S9305, 1:10,000 dilution).

### DNA isolation, deep sequencing, and read preprocessing and mapping

Total genomic and nucleolar DNA was isolated with Quick-DNA Universal Kit (Zymo Research, CA, USA). Libraries were constructed using Illumina’s TruSeq DNA PCR-free Library Preparation kit (350 bp) and fragments were size-selected by sample-purifying beads. 150bp paired-end sequencing was performed using Illumina HiSeq X Ten sequencing system using the HiSeq X Reagent Kit V2. 206.6 and 186.6 million reads for first replicate were obtained from genomic and nucleolar samples, respectively. >96% and >94%, respectively, of these were mappable. For subsequent samples, the subsampling analysis suggested that 50 million reads were sufficient (Supplemental Figure 3). For more details on sequencing, please see the files associated with the data at data.4dnucleome.org.

Sequencing reads were trimmed with cutadapt (version 1.16) (Martin 2011) before being aligned to the mouse genome (mm10) using Bowtie2 v2.1.0 with the standard default settings (Langmead and Salzberg 2012). Alignments with mapping quality score greater than 20 were kept (samtools v1.3), and duplicated reads were removed using picard tools v1.96 (hNps://broadinsUtute.github.io/picard/). To visualize the mapped reads, the bigwig files were generated using deepTools2 (Ramírez et al. 2016).

### NAD identification and annotation

To facilitate the analysis of this type of data, we implemented a common workflow for NAD-seq data analysis as a Bioconductor package called *NADfinder*.

To make it easy to use, all the functionalities for analyzing data with multiple biological replicates have been wrapped into four functions: *tileCount, log2se, smoothRatiosByChromosome* and *callPeaks*. An overview of the NADfinder workflow is depicted in Supplemental Figure S4A. Briefly, summarized counts for each moving window (default 50kb) with a step size (default 1 kb) are computed for nucleolar and genomic samples using the *tileCount* function. Log2 ratios of counts or CPM are computed between nucleolar and corresponding genomic samples for each window using the *log2se* function. Chromosome-level baseline correction and smoothing of Log2 ratios are performed using the function *smoothRatiosByChromosome*. Peaks/NADs across biological replicates are called, filtered and merged using *callPeaks*. We used default setting of *NADfinder* to analyze our data except the following parameter settings: lfc=log2(1.5), countFilter = 200 and cutoffAdjPvalue = 10^-3^ (10^-2^ for chrX, where the signal to noise ratio is less than other chromosomes).

Nucleotide-level overlap analysis of NADs and other genomic features such as LADs, H3K9me3 and H3K27me3 were performed using *GenomicRanges* (Lawrence et al. 2013). Specifically, the functions *intersect* and *setdiff* were used to obtain the overlapping nucleotides between two peak Granges, and the nucleotides in only one of the peak Granges, respectively. In addition, GO enrichment analysis of different classes of NADs were performed using *ChIPpeakAnno* (Zhu 2013; Zhu et al. 2010).

### DNA-FISH probes

The bacterial artificial chromosomes (BACs) were obtained from the BACPAC Resource Center of Children’s Hospital Oakland Research Institute (Oakland, CA). DNA was isolated using BAC mini DNA prep Kit (Zymo Research, CA). BAC probes were and labeled using the Bioprime Labeling Kit (Invitrogen Eugene, OR USA).

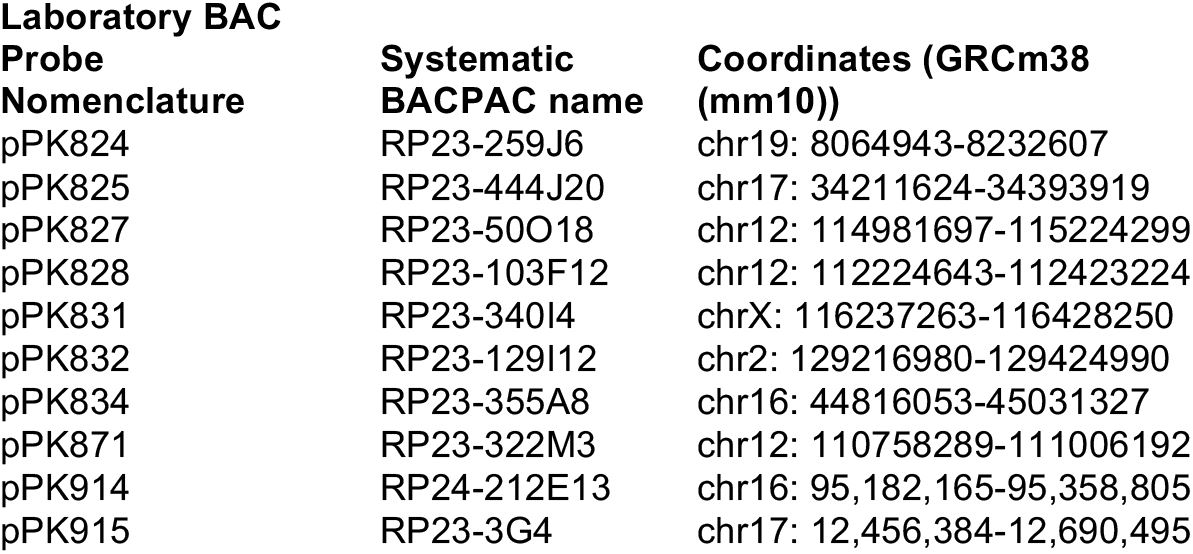

### 3-D DNA FISH/ immunocytochemistry and microscopy

Immunofluorescence microscopy analysis of DNA-FISH/immunocytochemistry-labeled cells was carried out as previously described (Smith et al. 2014; Matheson and Kaufman 2017). Briefly, cells were grown on sterile coverslips until ~70% confluent. For mES cells, serum was added to 2i medium (10% final) prior to seeding them on 0.1% gelatin-coated coverslips (10 hours). Cells were rinsed in PBS, placed in a coplin jar just long enough to rinse with 10 ml of CSK buffer (CSK Stock: 50 ml 0.1 M Pipes-NaOH pH 7.8; 50 ml 1M NaCl; 1.5 ml 1M MgCl_2_; 51.35 g sucrose. Total volume was brought to 500 ml with H2O, filter sterilized and stored at 4°C). Next, cover slips were transferred into a second coplin jar containing Triton-CSK buffer for 5 min (9 ml CSK buffer + 0.5 ml 10% TritonX-100 solution (Roche) + 0.5 ml 200 mM VRC (vanadyl ribonucleoside complex; NEB). After 5 min on ice, the samples were transferred into a third jar containing 10 ml of 4% Paraformaldehyde in 1X PBS for 10 min at RT. Samples were rinsed with RT 70% EtOH 3x, and transferred into a coplin jar containing 70% EtOH for 10 min at RT.

After 10 min of incubation in 70% EtOH, the coverslips were incubated in RNA removal solution (10 ml 70% EtOH + 200 μl 10M NaOH) for 5 min at RT. Coverslips were then washed 3 times with 70% EtOH, and then incubated for 5 min in 70% EtOH and in 100% EtOH for another 5 min. The coverslips were then briefly air dried prior to denaturation at 80^0^C for 2.5 min in a buffer containing 7 ml Formamide, 2 ml H_2_O, 1 ml 20 x SSC; pH adjusted between 7.1 and 7.4 with HCl, pre-heated in coplin jar to 80^0^C. After denaturation, coverslips were incubated for 5 min in ice-cold 70% EtOH and then for 5 min in ice-cold 100% EtOH.

DNA hybridization mixtures contained: 3 μl Cot-1 DNA (1 μg/μl), 2 μl salmon sperm (Invitrogen, D9156) and added tRNA (10 mg/ml), and 1-5 μl of Bioprime labelled FISH DNA Probe (100 ng/coverslip). These mixtures were assembled in 1.5 ml eppendorf tubes on ice and placed into a Speed-Vac for 20 minutes – 1 hour until the liquid completely dried. The DNA was resuspended with 10 μl formamide (Sigma-Aldrich 221198), vortexed and spun at 16,000 x g for 1 min. The samples were heated to 80°C for 10 min and 10 μl of hybridization buffer was added to the tube. This hybridization cocktail was transferred to parafilm on a glass support. The coverslips were briefly air dried, inverted onto the hybridization cocktail and covered with parafilm. Coverslips were then incubated O/N at 37^0^C in a humidified chamber. After incubation, samples were washed first with 5 ml formamide + 5 ml 4X SSC for 20 min at 37°C in a humid chamber; then with 2X SSC for 20 min in 37° C humid chamber; and with 1X SSC, at RT in a shaker for 20 min. Coverslips were incubated with 4X SSC at RT for 1 min. Fluorescently-labeled Streptavidin (594 or 488) was diluted 1:500 in 4X SSC / 1%BSA and coverslips were incubated with this at 37°C for 1 hour. The incubation was followed by three washes perfomed in the dark: 4X SSC at RT in a shaker for 10 min; 4X SSC / 0.1% Triton at RT in a shaker for 10 min; and with 4X SSC at RT in a shaker for 10 min.

Coverslips were then incubated in PBS for 5 min. For immunocytochemistry, samples were blocked in 1X PBS/ 1% BSA for 15-30 min at RT. Primary antibodies were diluted in blocking solution and samples were incubated O/N at 4^0^C. Samples were washed with PBS and incubated with secondary antibodies for 1 h at 37 ^0^C. Coverslips were washed with PBS, 1X PBS / 0.1% TritonX-100, and 1xPBS at RT on a shaker, 10 min each. Finally, the coverslips were rinsed twice with PBS and one time with water, and mounted using Prolong Gold antifade reagent (Invitrogen, cat#P36934).

Images were acquired with a Zeiss Axiovert 200M, a Perkin Elmer Ultraview spinning disc microscope and Hamamatsu ORCA-ER camera. (100x NA1.4 Plan-Apocromat Oil objective). DNA-FISH probes were counted through Z-stacks manually, and were scored as “associated” when no detectable gap between nucleolar marker and labeled probe was detected throughout the Z-stack. Each probe was analyzed in at least three biological replicates, with about 100 alleles counted per each replicate. For images, fluorescence range intensity was adjusted identically for each series of panels. *Z* stacks are shown as 2D maximum projections or as a single optical section (MetaMorph, Molecular Devices). All statistical analysis was done using GraphPad Prism software.

Fluorescence intensities were quantified from the sum of Z-stacks using an adaptation of previously described methods (Delaval et al. 2011). Intensities in concentric circles of 408 μm^2^ (inner area) and 816 μm^2^ (outer area) were measured. Inner area circles were placed around nuclei so that the borders of the nucleus did not go beyond the area. Nuclear signal intensity was calculated using formula: I=E5-((E6-E5)*(C5/(C6-C5)), where I is nuclear intensity; E5 is an integrated signal intensity of inner circle; E6 is an integrated signal intensity of outer circle; C5 is the area of inner circle, C6 is the area of outer circle. In this manner, local background was calculated as the signal in the outer area minus the signal in the inner area, and the signal in each nucleus was the inner signal minus the local background.

## Supporting information

## Data access

All sequencing data and quality control information for individual samples is publically available at the NIH 4DN Data Portal (data.4dnucleome.org) under accession numbers 4DNES15QV1OO, 4DNES8D85R8H, 4DNESR3K7VRN, 4DNES4U7AY3P, 4DNESNEBF9W4, 4DNES7SVJXAI. See Supplemental Table S1 for details.

## Acknowledgements

We thank Thoru Pederson for review of this manuscript. Research reported in this publication was supported by the National Institutes of Health (U01 DA040588 to P.D.K.) as part of the 4DN consortium.

---

**Supplemental Figure S1.**
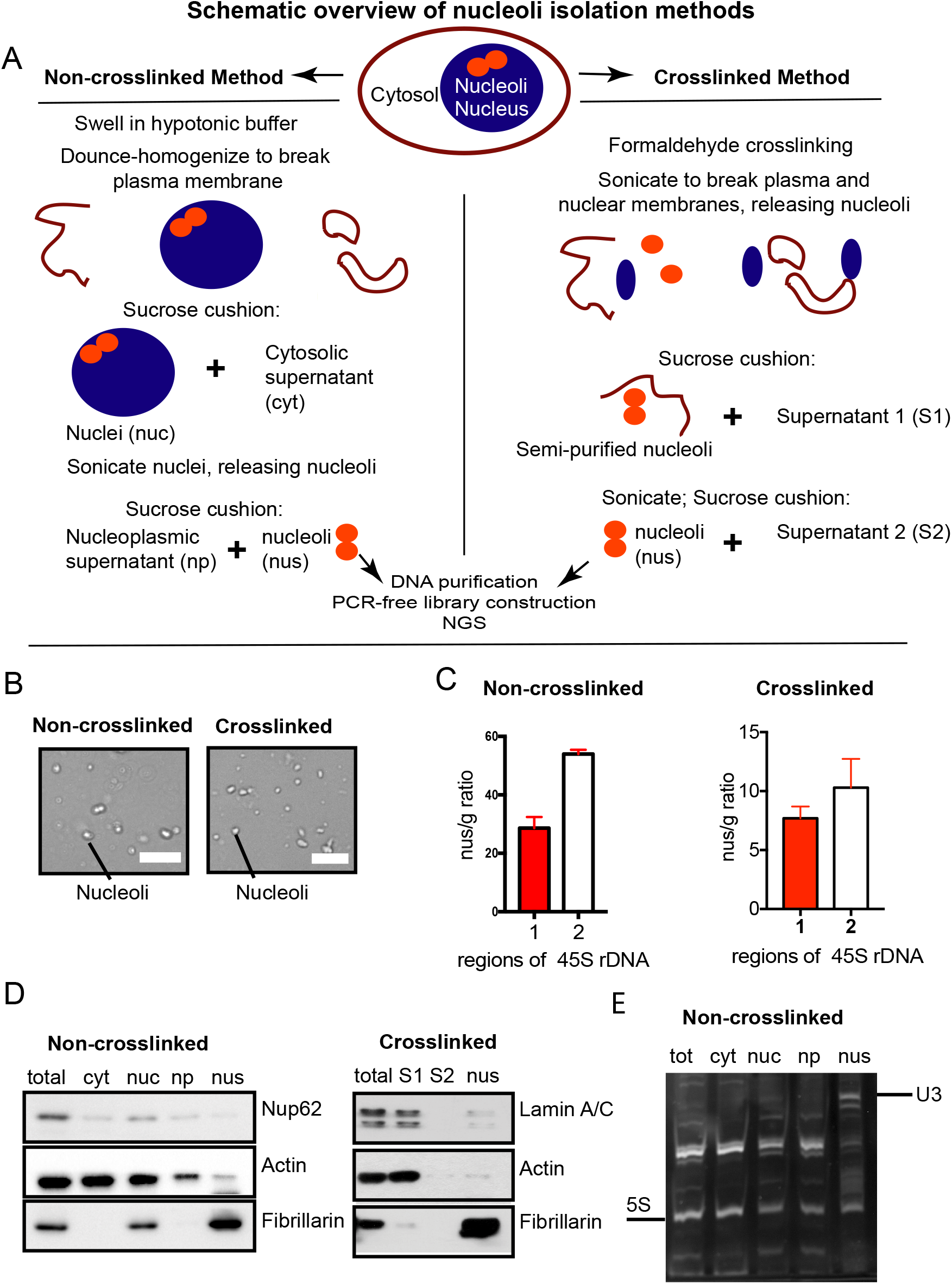
Isolation and characterization of purified nucleoli. A. Schematic representation of crosslinked and non-crosslinked methods used for isolation of nucleoli. The abbreviations of the fractions analyzed by immunoblotting in panel D are indicated.
B. Phase microscopy images of nucleoli purified via crosslinked (experiment #24, see Supplemental Table S1) and non-crosslinked (experiment #29) methods (20x magnification, scale bar 10 μm).
C. qPCR analyses of rDNA enrichment in purified nucleoli compared to genomic DNA. We observed 30-50-fold rDNA enrichment in purified non-crosslinked nucleoli (experiment #33), depending on the primer set used. An 8-13-fold enrichment was observed in crosslinked samples (mean values for experiments #24, 26, 28).
D. Immunoblot analyses of crosslinked and non-crosslinked preparations. Fibrillarin was used as a nucleolar (nus) marker. Nuclear periphery proteins (porin Nup62 and LaminA/C) were present in total (tot) and crude nuclear (nuc, S1) extracts, but depleted from nucleolar (nus) fractions. Cytoskeletal protein actin was enriched in total and crude cytosolic (cyt, S1) fractions, but not in nucleoplasmic (np) or nucleolar (nus) fractions.
E. Analysis of RNAs from fractions obtained during a non-crosslinked preparation. The nucleolar-specific small RNA U3 (Politz et al. 2009) as well as the ribosomal 5S species are illustrated.

**Supplemental Figure S2.**
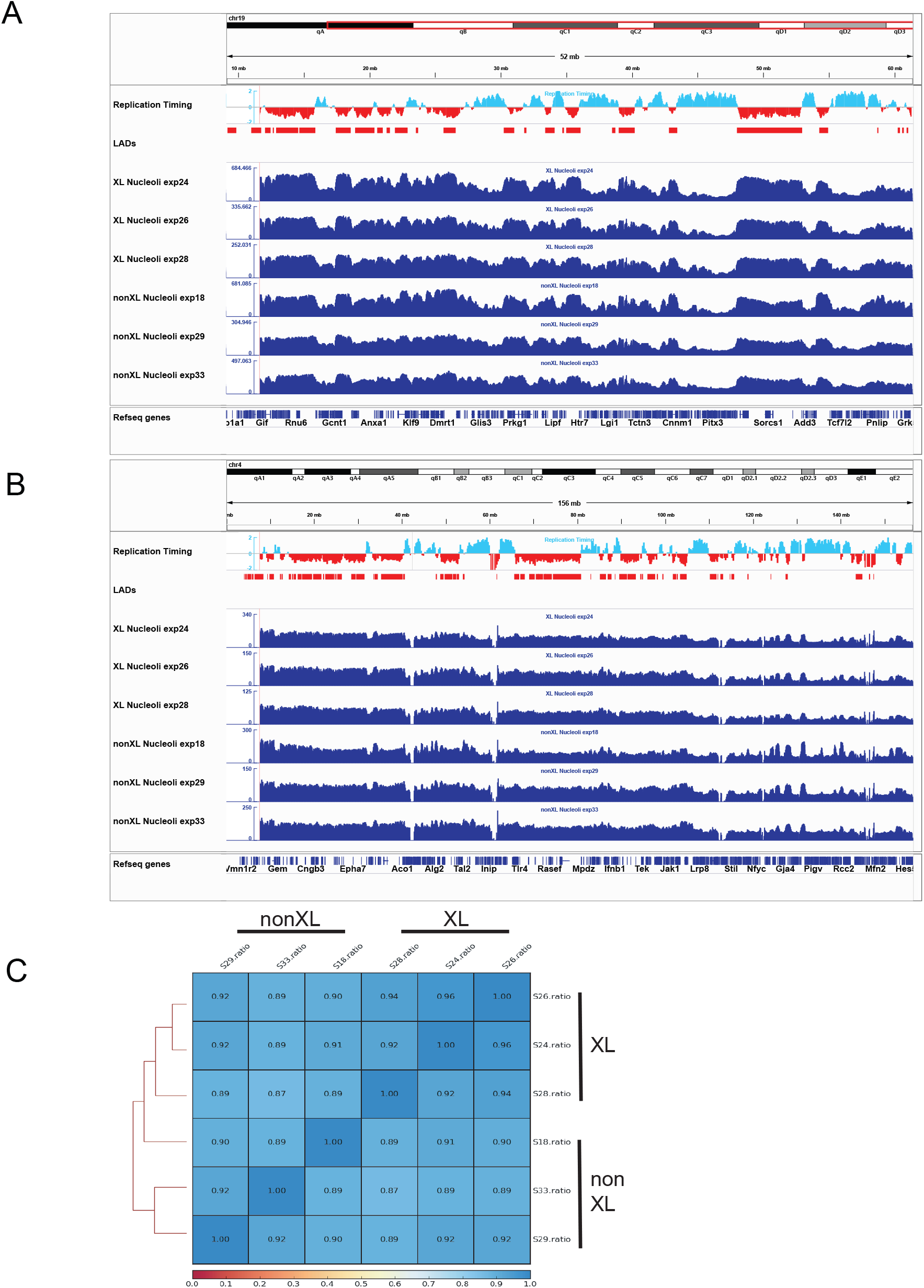
Analysis of raw data from biological replicate samples. In addition to DNA replication timing (early replicating regions in cyan, late in red, (Hiratani et al. 2010)) and LAD (Peric-Hupkes et al. 2010) data from MEFs at the top, we display the raw sequencing reads from purified nucleoli from each of six biological replicate experiments. Three samples were made using the crosslinked protocol (XL), and three were non-crosslinked (nonXL). A. Chromosome 19.
B. Chromosome 4.
C. Pearson correlations of log2 (Nucleolus/Genomic) ratio of all 6 experimental samples were calculated. Crosslinked (XL) and non-crosslinked (nonXL) samples are indicated.

**Supplemental Figure S3.**
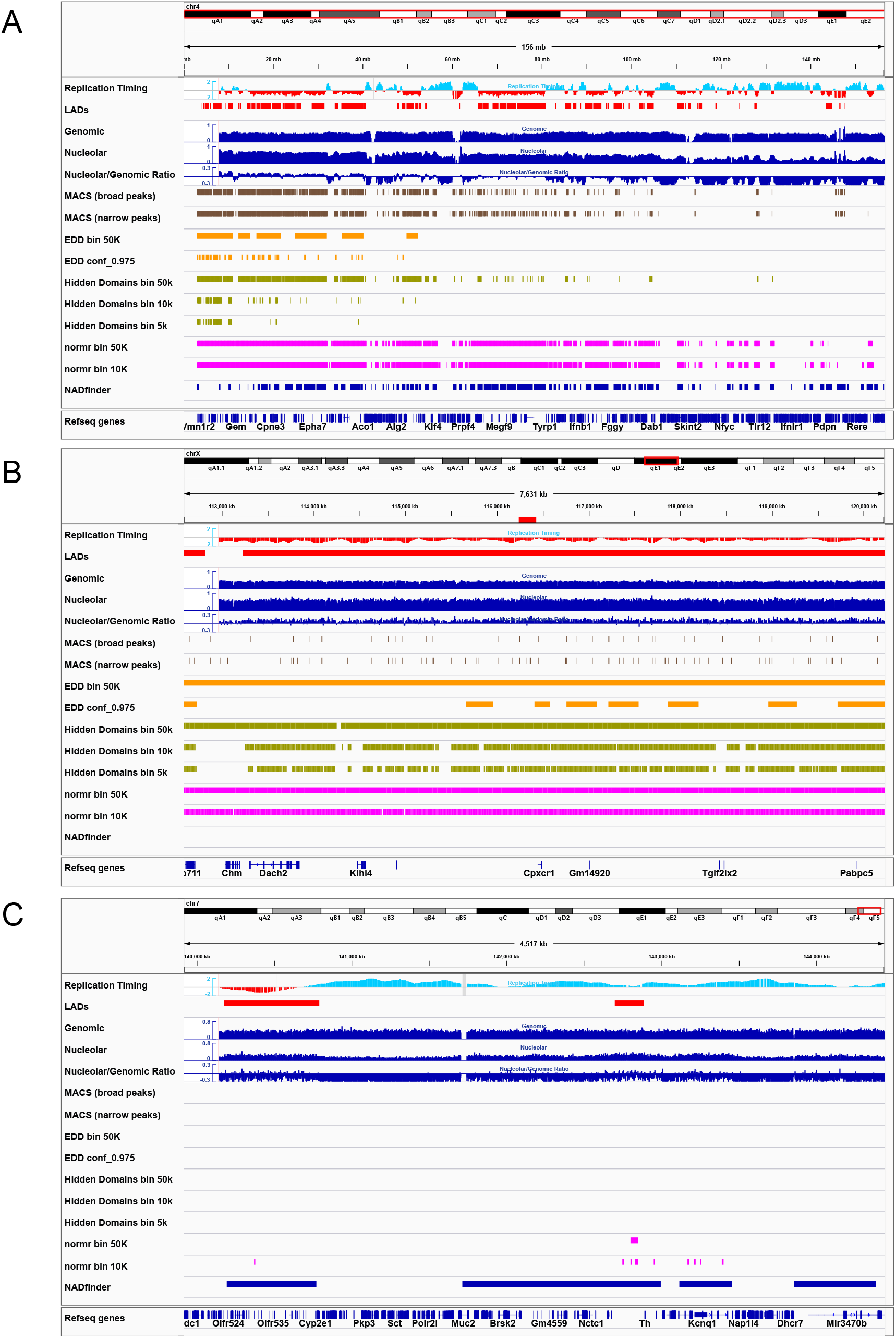
Additional comparisons of different bioinformatic analyses of NAD-seq data. A. Chromosome 4 is shown in its entirety. In addition to the tracks shown in Figure 1B-C, we compared additional settings for each bioinformatics program tested. For MACS, we compared the broad peak, and narrow peak settings (Zhang et al. 2008; Feng et al. 2012). For EDD, we compared 50 Kb binning with and without filtering at a 0.975 confidence threshold (Lund et al. 2014). For Hidden Domains, we compared binning at 5, 10, and 50 Kb (Starmer and Magnuson 2016). For normr, we compared binning at 10 and 50 Kb (Kinkley et al. 2016; Helmuth et al. 2016). As observed for chr5 (Fig. 1C), unlike MACS and EDD, NADfinder was capable of identifying peaks in the region distal from the centromere (on the right).
B. As in panel A, showing a region of chrXqE. Note the flat level of nucleolar enrichment across this region. The red bar at the top is probe pPK831, a BAC that was observed to be negative for nucleolar association in DNA-FISH experiments (see Fig. 2). This region was correctly called negative by NADfinder, but incorrectly identified as positive by EDD, Hidden Domains, and normr.
C. As in panel A, showing a distal telomere-proximal region of chr7, including the Kcnq1 locus. This locus has previously been shown to associate with nucleoli (Pandey et al. 2008; Fedoriw et al. 2012a). Of the bioinformatics programs used, only NADfinder correctly called this locus as a peak.

**Supplemental Figure S4.**
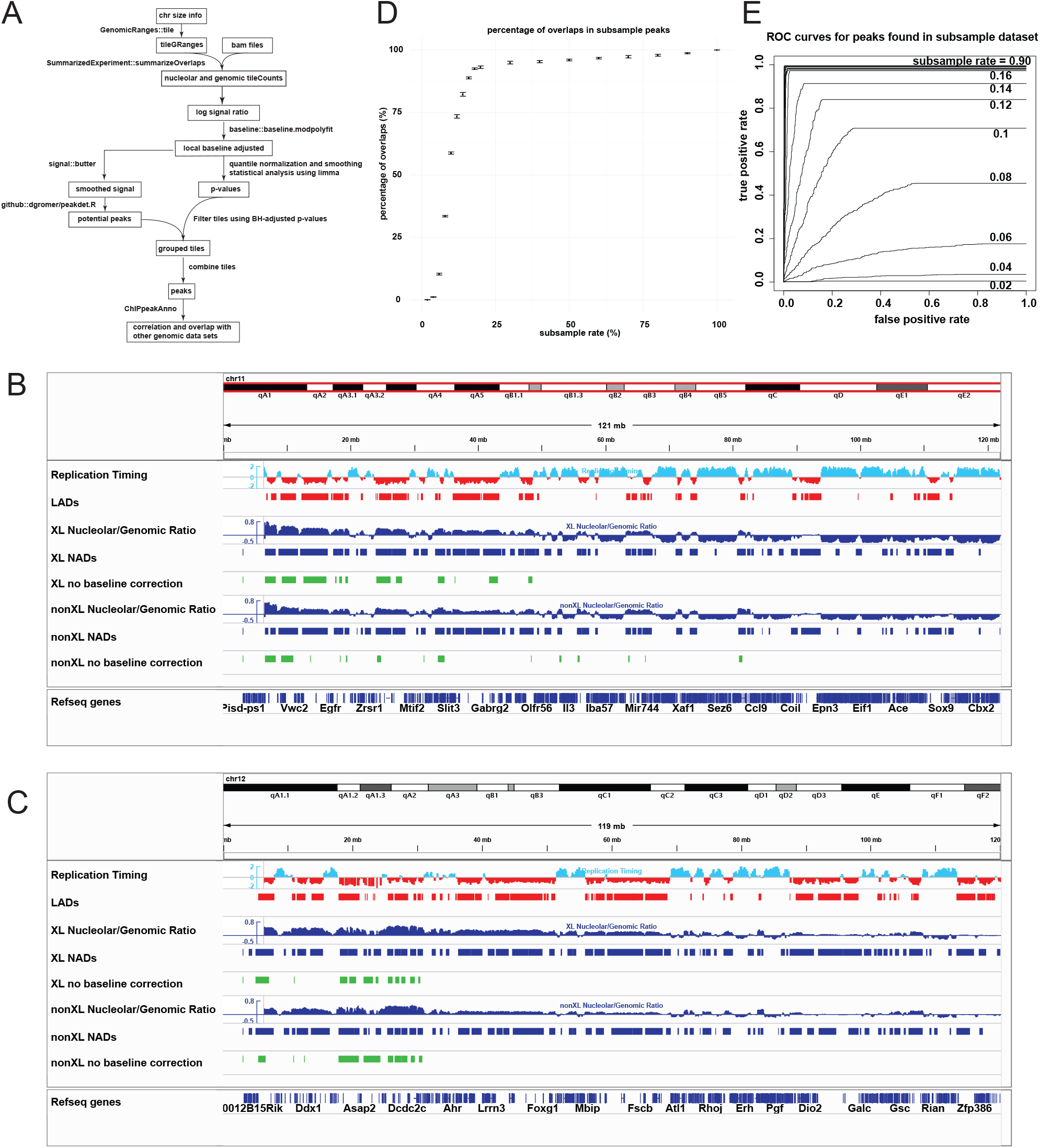
Testing baseline correction and sequencing depth requirements for *NADfinder*. A. Workflow of NAD-seq analysis using *NADFinder* and *ChIPpeakAnno*.
B. Chromosome-level baseline correction is necessary for optimal peak calling by *NADfinder*, shown for chr11. Without baseline correction, peaks at the distal region of the chr11 are missed. Replication timing and LAD peak data from MEFs are shown as in Fig. 1. Blue tracks show nucleolar/genomic ratios from crosslinked (XL) and noncrosslinked (nonXL) preparations, with *NADfinder* peaks also in blue. Green tracks show peaks called by *NADfinder* without baseline correction.
C. As in panel B, chr12 is shown. Same as chr11, without baseline correction, peaks at the distal region of the chr12 are missed.
D. Subsampling analysis with peak calling at 5% false discovery rate. In this experiment, about 200 million reads per library were obtained for 2 pairs of genomic and nucleolar samples. This analysis suggests that 25% of subsampling rate resulting in about 50 million reads, would identify similar sets of peaks at 5% discovery rate as the whole dataset, i.e. 200 million reads.
E. Receiver operating characteristic curves (ROC) show that a 20% subsampling rate results in similar ROC as observed using the entire 200 million read dataset in panel D.

**Supplemental Figure S5.**
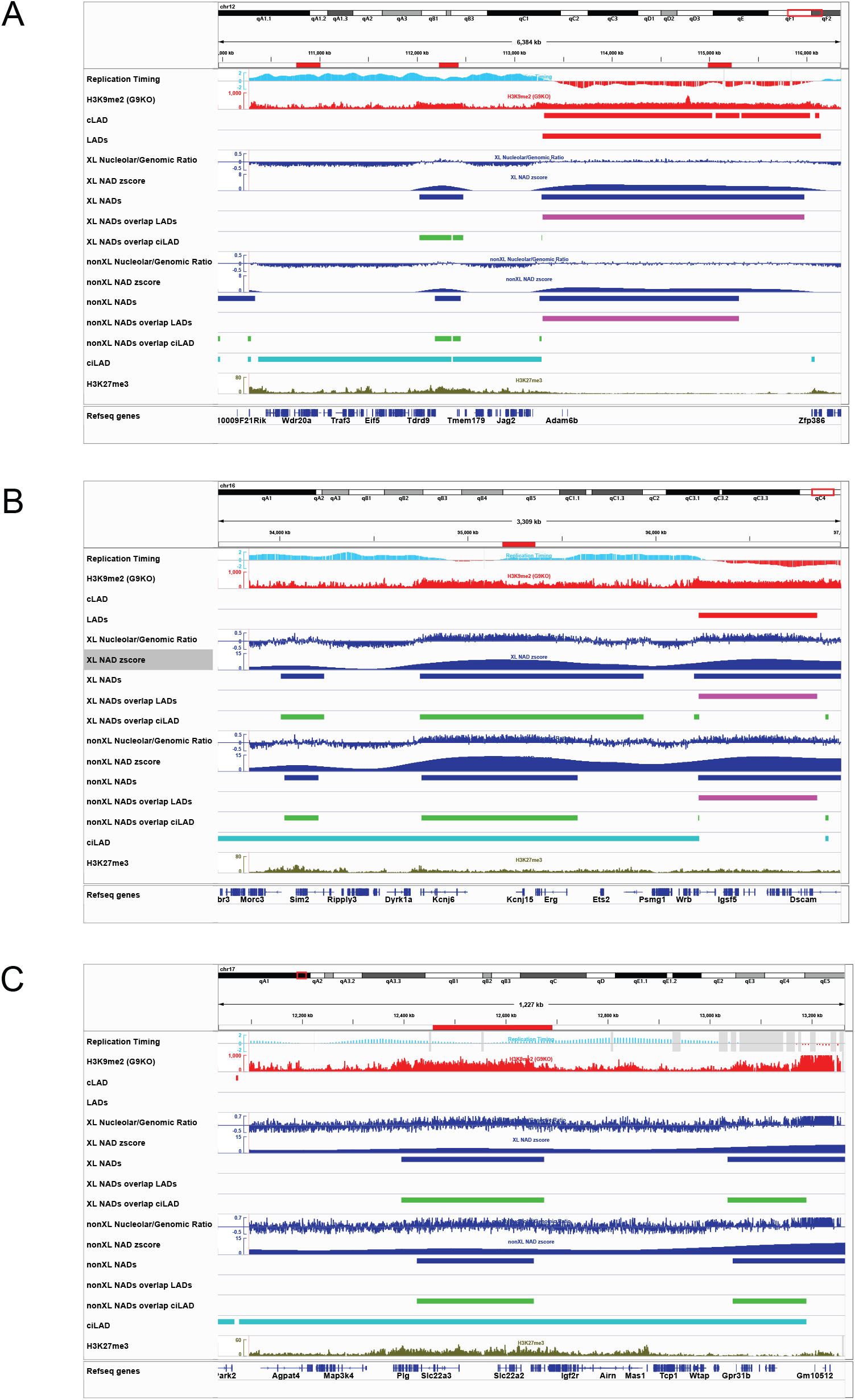
Genomic locations of BACs used for FISH experiments. BAC locations are shown as red horizontal bars at the top of each graph. A. The same IGV session used in Figure 6 is used here to show a region of the chr12qF locus that contains three BAC probes used in these studies. Left to right, pPK871 (negative control, no NAD enrichment), pPK828 (Type II NAD, overlapping ciLAD but not LAD regions), and pPK827 (Type I NAD, overlapping LAD but not ciLAD regions).
B. pPK914. This Type II NAD contains ion channel genes (Kcnj6, Kcnj15) as well as developmentally regulated Ets-family transcription factors (Erg, Ets2). The Type II NAD to the left overlaps the Sim2 gene encoding a developmentally-regulated, single-minded family bHLH transcription factor.
C. pPK915. This Type II NAD contains a cluster of cation transporter genes (Slc22a1,2,3).

**Supplemental Figure S6.**
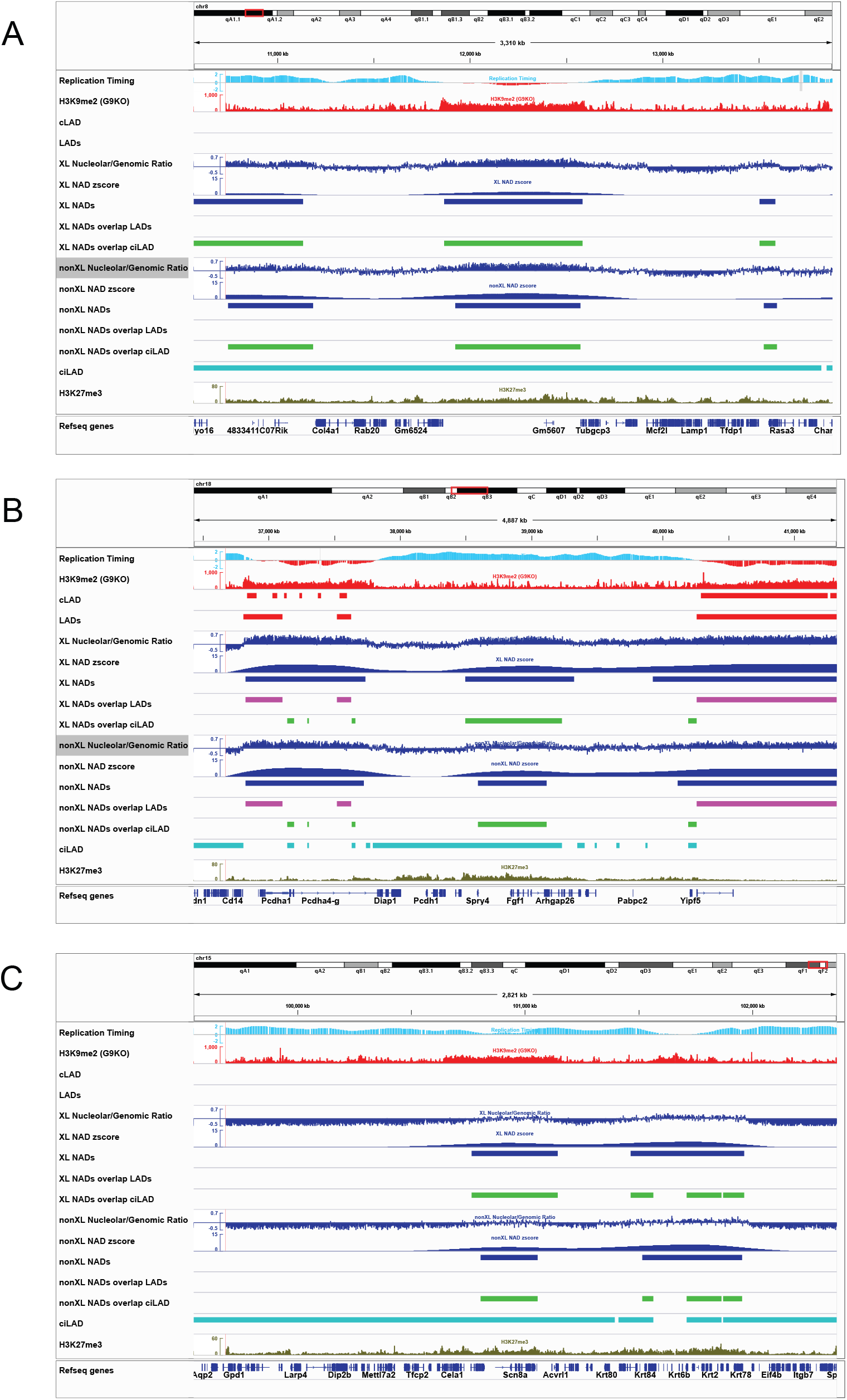
Examples of Type II NADs carrying genes with enriched GO annotations. A. The same IGV session used in Figure 6 is used here to show a region including the NAD encompassing the gene encoding developmental transcription factor Sox1 (which is very small and located within the Gm5607 locus).
B. This Type II NAD contains the gene encoding the Fibroblast growth factor 1 (Fgf1) gene.
C. There are two Type II NADs shown here. On the left is one containing the Scn8a gene encoding a sodium channel. On the right is a NAD containing a cluster of keratin (Krt) genes.

Supplemental Table 1

Sequencing depth/mapping statistics for replicate experiments

Supplemental Table 2

GO term analyses of Type II NADs, showing annotations with q< 0.05

